# Millisecond-scale, single-neuron credit assignment in a songbird

**DOI:** 10.64898/2026.07.31.740148

**Authors:** Josefa R. Scherrer, Jacob M. White, Alison N. D. Plump, Michale S. Fee

## Abstract

Learning an adaptive behavior requires identifying which actions, in which contexts, lead to particular outcomes. This problem of credit assignment is fundamental to both biological and artificial learners^1–3^. Songbird vocal learning presents a particularly demanding credit assignment problem: singing is controlled by millisecond-precise activity of thousands of motor neurons^4–7^, but song quality is encoded by a diffuse, delayed dopamine signal with more than an order of magnitude less temporal precision^8–12^. It is unknown how precisely the songbird brain can drive changes in specific premotor neurons at precise times to improve song performance. Here we show that the cortico-basal ganglia circuit thought to underlie songbird vocal learning can learn with millisecond-scale temporal precision and single-neuron spatial precision. We find that playing disruptive auditory feedback contingent on the activity of a targeted neuron in a vocal variability-generating premotor nucleus at one time in the song causes adaptive changes in the activity of the targeted neuron with 3.2 ms temporal precision. Learned changes in firing rate are not observed in uncorrelated neighboring neurons, thus revealing single-neuron spatial precision. These findings challenge the prevailing view of dopamine-mediated reinforcement as slow and imprecise^13–15^, and redefine our understanding of the limits of credit assignment in the brain.

## Main

Complex motor behaviors, like speech and musical performance, rely on precisely timed sequences of actions that must be learned. It is thought that many behaviors like these are acquired through a trial-and-error process of reinforcement learning (RL). However, while motor behaviors require the precise coordination of many neurons, acting on multiple muscles on a millisecond timescale^16^, dopaminergic reinforcement signals in the brain are thought to be low-dimensional, slow, and delayed^17^. It remains unclear how the brain utilizes such a relatively uninstructive reinforcement signal to craft the precise, moment-by-moment dynamics of complex motor behavior. In particular, how is credit for a global dopaminergic fluctuation assigned to the specific neurons and precise timepoints that caused it? Here, we set out to address this question in the context of song imitation by the zebra finch (*Taeniopygia guttata*).

The courtship song of the zebra finch is an exquisitely precise motor behavior driven by equally precise neural activity. Adult male zebra finches sing a stereotyped motif of 3-7 song syllables that each contain acoustic features modulated on timescales as short as 5 ms^18,19^. Song production is subserved by the specialized song motor pathway (SMP), which comprises premotor nucleus HVC (used as a proper noun) and motor cortex analog RA (robust nucleus of the arcopallium) (Fig. 1a). During singing, the majority of active HVC neurons each fire at a single moment in time and as a population form a continuous, sparse sequence with sub-millisecond temporal precision^20,21^. These HVC neurons drive the ~8000 downstream RA neurons to burst in stereotyped sequences with similar temporal precision^4–6^. Then, via projections to lower motor neurons in the brainstem, RA drives the actions of the ~8 muscles in the vocal organ^22,23^.

**Figure 1:**
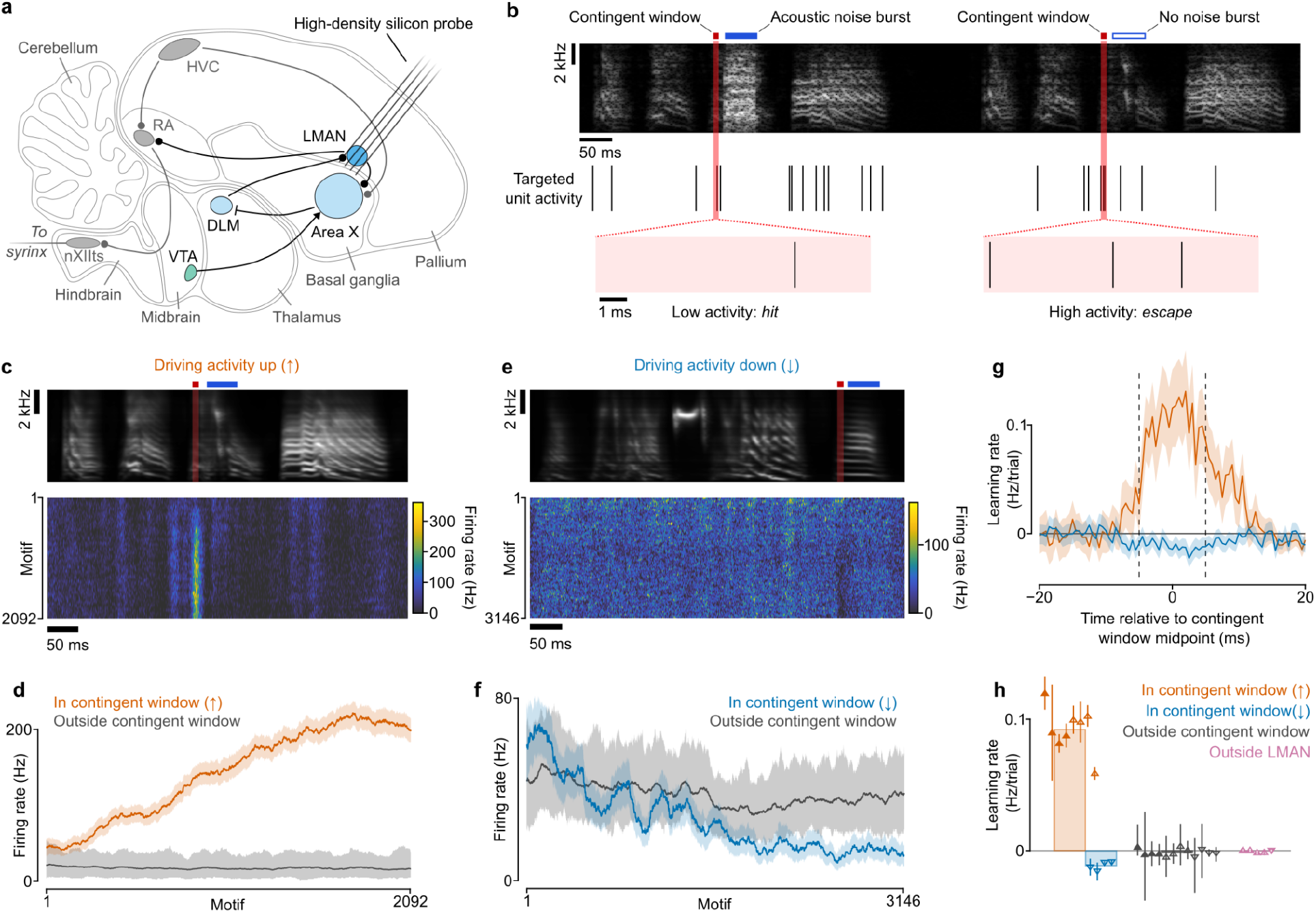
LMAN activity changes in response to neuron-conditional auditory feedback (nCAF). **a**, Neuropixels 2.0 probes were implanted in premotor nucleus LMAN and used to record neural activity and perform closed-loop feedback. **b**, During nCAF, the activity of a targeted neuron is measured within a narrow time window (the *contingent window*, red) at a consistent time in the song. On each motif, a brief acoustic noise burst (*disruptive auditory feedback*, DAF; blue) is played to the bird contingent on the number of spikes the targeted neuron generates within the contingent window on that motif. In this example, DAF is played (hit) when the number of spikes is below average (left) and DAF is withheld (escape) when the number is above average (right). **c**, Results of an nCAF experiment on a targeted single neuron where DAF was played on low activity in a 10 ms contingent window. (Top panel) Song motif. (Bottom panel) Each row shows the song-aligned firing rate of the targeted single neuron during each song motif, plotted in the order in which motifs were sung throughout the day. Firing rates are smoothed with a two-dimensional Gaussian kernel with widths of 0.4 ms and 20 motifs. **d**, (orange) Firing rate within the contingent window over the course of the day in panel c. (gray) Firing rate outside the contingent window. (Shading is 95% CI; firing rates are calculated over a 100 motif sliding block. **e**,**f** Same as panels c and d, but for a targeted neuron where DAF was played when activity was higher than average. **g**, Learning rate (*i*.*e*., slope of firing rate versus motif number) at different times relative to the contingent window (vertical dashed lines) for the targeted neurons in panel c (orange) and panel e (blue, see Methods, envelope is 95% CI). **h**, Summary plot showing learning rates for nCAF experiments. All within-LMAN datapoints driving up (orange; n=4 single units plus 4 multi-unit signals in 7 birds) or down (blue; n=4 units in 3 birds) exhibit significant learning (CI does not include zero). Shaded triangles are single units, hollow triangles are multiunit signals. Direction of triangle indicates DAF contingency direction. Error bars for each point are 95% CI of the mean.

Remarkably, this ultra-precise vocal performance is learned by trial and error. Male zebra finches learn to imitate the song of a tutor bird through weeks of practice. This learning process requires a specialized cortico-basal ganglia circuit, the anterior forebrain pathway (AFP)^24–26^. The AFP is composed of basal ganglia nucleus Area X, thalamic nucleus DLM (medial nucleus of the dorsolateral thalamus), and pallial (cortex-like) nucleus LMAN (lateral magnocellular nucleus of the anterior nidopallium), organized in recurrent, topographic loops^27^ (Fig. 1a). Vocal exploration is driven by LMAN, whose highly variable firing patterns perturb the song via topographic projections to RA^27^. The quality of a given song variation is evaluated in auditory cortex (pallium)^28^ and conveyed to Area X by a dopaminergic reward-prediction error (RPE) signal from neurons in the ventral tegmental area (VTA)^8–11^. Notably, as in other species, this dopaminergic RPE signal appears to be low-dimensional and fluctuate at a slower timescale, and with a substantial delay, relative to the vocal errors it encodes: observed transient changes in VTA activity are on the order of 100 ms long, and follow song changes with a ~50 ms latency^8,9^.

Previous studies have shown that birds can learn rapidly in response to presentation of disruptive auditory feedback (DAF) that is contingent on song features. For example, playing a burst of acoustic white noise on song motifs where the pitch at a particular time is lower than average will cause the song pitch at that time to increase gradually over the course of several hundred song renditions (pitch-conditional auditory feedback or pCAF^29–31^). Critically, this learned change initially requires LMAN in order to be expressed^30,31^, suggesting that song changes are driven first by changes in LMAN activity.

This finding led to a model of song learning in which medium spiny neurons (MSNs) in Area X detect which patterns of LMAN activity, at each time in song, are correlated with subsequent dopamine increases. Then, via a multisynaptic pathway including DLM, the MSNs bias those same LMAN neurons to be more active at the same time in the song on subsequent song renditions^32^. Notably, this mechanism could allow for highly spatiotemporally precise learning, despite sluggish dopamine signals. This model has been generally supported^30,33,34^, but its fundamental prediction—that the activity of LMAN neurons becomes biased towards patterns associated with improved vocal performance—has never been directly tested.

To test this model, we designed a neural feedback system (neuron-conditional auditory feedback or nCAF) in which we measured the activity of a targeted LMAN neuron at a precise time in the song within a short time window (the *contingent window*). We then delivered disruptive auditory feedback (DAF) conditional on the number of spikes in the contingent window on each song motif and found that nCAF drove precisely timed learned changes in the activity of individual LMAN neurons. Our results support a specific model of basal-ganglia-mediated learning capable of solving the credit assignment problem with single-milliseconds temporal precision and single-neuron spatial precision.

### Targeted learning driven by closed-loop auditory feedback

Our nCAF neurofeedback system is based on chronic Neuropixels 2.0 recordings using a custom microdrive implant^35^ (Fig. 1a). nCAF was carried out over the course of an entire day, during which we recorded the activity of a population of LMAN neurons. In a first set of experiments, we played DAF (white noise burst, see Methods) on motifs when the targeted neuron generated fewer than the median number of spikes within a 10 ms contingent window (see Methods). DAF was played with 5-15 ms latency following the end of the contingent window (see Methods). We found that in response, the activity of the targeted neuron reliably increased within the contingent window over a single day (Fig. 1c,d, example single neuron; Fig. 1h, 7 birds, n=4 single units plus 4 multiunit signals; learning rate = (9.2 ± 0.6)×10^−2^ Hz/motif, mean ± s.e.m.; t_7_=15, p=8.0×10^−7^, one-sided t-test; Extended Data Fig. 1). Learned changes were largely confined to the contingent window (Fig. 1g), with an approximate width of 12 ms (full width at half maximum, comparable to the width of the contingent window; equivalent to a Gaussian with s.d. of 5.1 ms). Firing rates did not significantly change at times in the song more than 10 ms from the edges of the contingent window (Fig. 1d,h, n=8 units in 7 birds, learning rate = (−0.92 ± 1.03)×10^−3^ Hz/motif, mean ± s.e.m.; t_7_=-0.89, p=0.40, two-sided t-test).

We next asked whether the direction of learning could be reversed by playing DAF when the targeted neuron had higher than median activity within the contingent window. Indeed, in a new set of experiments, the activity of the targeted neuron decreased within the contingent window over the course of the day (Fig. 1e,f, example unit; Fig. 1h, 3 birds, n=4 units, learning rate = (−1.3 ± 0.2)×10^−2^ Hz/motif, mean ± s.e.m.; t_3_=-8.0, p=2.0×10^−3^, one-sided t-test) while remaining unaffected at other times in song (Fig, 1e-h, 3 birds, n=4 units, learning rate = (−2.9 ± 1.4)×10^−3^ Hz/motif, mean ± s.e.m.; t_3_=-2.1, p=0.13, two-sided t-test).

We wondered whether the ability to learn in response to auditory feedback is a specialized property of LMAN, or is a general property of cortex-like (pallial) neurons. In contrast to LMAN neurons, nearby nidopallial neurons outside of LMAN (see Methods) did not learn when targeted with feedback in either contingency direction (Fig. 1h, n=4 single units and 1 multiunit signal in 3 birds, rate = (−3.6 ± 4.0)×10^−4^ Hz/motif; t_4_=-0.90, p=0.42, two-sided t-test).

### Learning is precise on a millisecond time scale

These findings suggest that song learning circuitry can alter neural activity in LMAN with a temporal precision comparable to the width of our contingent window (10 ms). However, in these measurements, the precision was limited by jitter in the targeting of the contingent window relative to song (± 2.5 ms range). To more accurately measure the temporal precision of learning, we performed an additional series of nCAF experiments using an even narrower contingent window (5 ms width) aligned more precisely to song timing (± 235 μs jitter, 95% CI, see Methods). To validate this increased temporal precision, we computed the correlation between spike count and DAF escape at each time in the song motif. We refer to this curve as the *correlation profile* (Fig. 2a). By fitting a Gaussian to the correlation profile for each targeted unit, we found that this more precise nCAF targeting induced correlations with temporal widths of 3.0 ± 0.9 ms (s.d. of Gaussian fit, mean ± s.d., n=15 units in 3 birds; Extended Data Fig. 2a-d). Note that the full width at half maximum of this correlation profile, 7.0 ± 2.2 ms (mean ± s.d.) is greater than the width of the contingent window, but is consistent with what we would expect based on the temporal autocorrelations of LMAN neurons (Extended Data Fig. 3).

**Figure 2:**
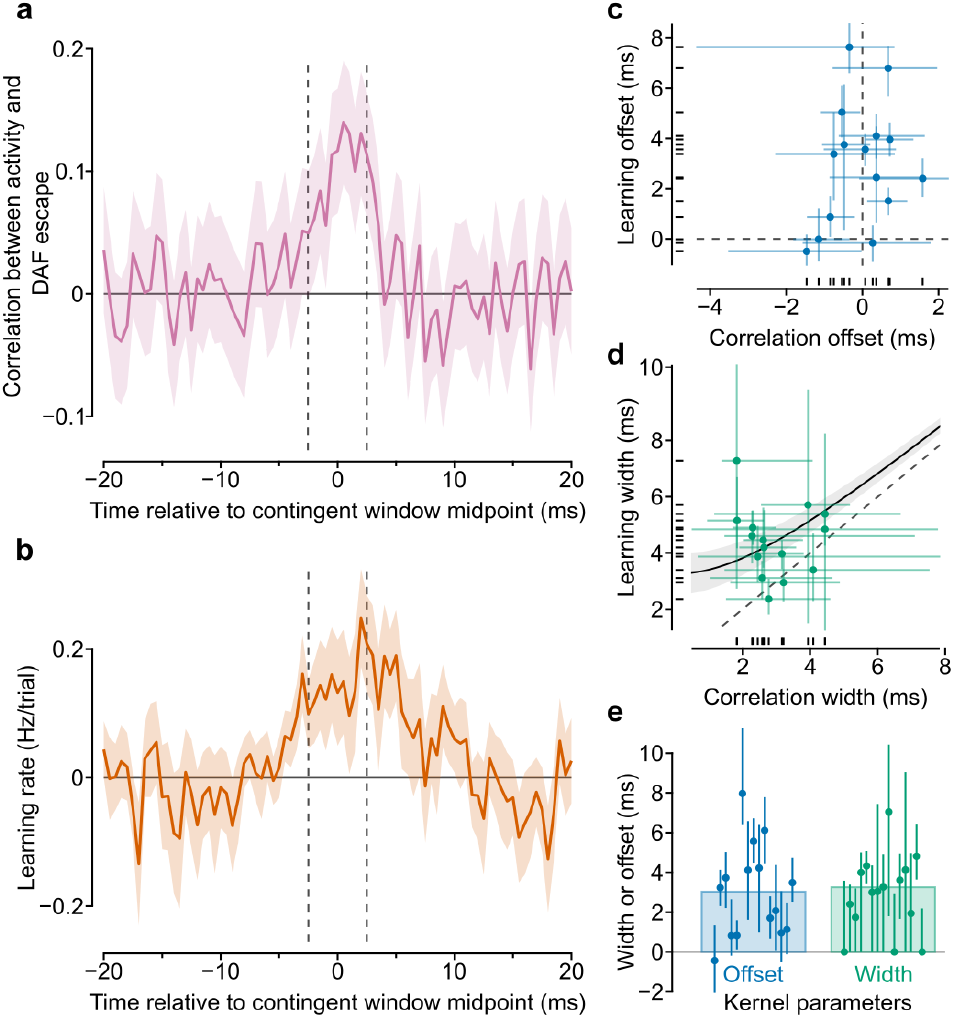
nCAF-driven learning is precise on a millisecond timescale. **a**, Temporal profile of correlation between activity of an example targeted neuron and DAF escape, imposed by the high-precision nCAF system. **b**, Temporal profile of learning rate for the same neuron after a day of learning. Envelopes are 95% CI. **c**,**d**, Gaussian functions are fit to the correlation profile and learning profile (from panels a and b) of each neuron on which nCAF is carried out, to extract the width and the temporal offset of these profiles (n=15 units in 3 birds). **c**, Offsets (Gaussian center) and **d**, widths (Gaussian standard deviation) of these fits (error bars are 95% CI on the fit parameters; axes are range bars; dashes along axes are marginal distributions). Curve in **d** is a hyperbola of best fit used to extract the underlying learning kernel width (shading is 95% CI). **e**, Offset and width (standard deviation) of the deconvolved learning kernel for each unit (error bars are 95% CI).

We next computed the rate at which the targeted unit increased its firing in the contingent window. This was computed at each time in the song motif to produce a curve that we refer to as the *learning profile*. We fit these curves for each targeted unit with Gaussians. The observed learning profiles had widths ranging from 2.4 to 7.3 ms, with a mean of 4.4 ± 1.2 ms (s.d of Gaussian fit, mean ± s.d., n=15 units in 3 birds, Fig 2d).

If the song learning circuit had perfect temporal precision, we would expect that the learning profile of each unit would exactly match its correlation profile. By contrast, if the circuit were highly imprecise, we would expect the learning profile to be much broader than the correlation profile. More generally, we can model the learning profile as the convolution of the correlation profile with an underlying Gaussian learning kernel. The width (s.d.) of the learning kernel is a measure of the precision of the system, with narrower widths representing higher precision. The offset (mean) of the kernel is a measure of how accurately the peak of the learning profile aligns with the peak of the correlation profile. We inferred the parameters of this learning kernel by deconvolving the Gaussian fits of the correlation profiles from the fits of the learning profiles, yielding an average offset of 3.0 ± 2.3 ms (mean ± s.d.) and average width of 3.2 ± 0.4 ms (mean ± s.d., Fig. 2e, n=15 units in 3 birds), thus demonstrating that the song learning circuitry can drive learned changes in LMAN activity with millisecond-scale temporal precision.

### Learned changes are partially driven by increased bursting

LMAN neurons exhibit burst events during singing that have been proposed to play an important role in learning^36–39^. We wondered whether the increase in spiking seen in our nCAF experiments was especially driven by an increase in burst events. We started by characterizing the prevalence of bursts in our recorded LMAN neurons at baseline (see Methods) and found that they occurred at a rate of 1.2 ± 0.2 per second during singing (mean ± s.d., n=13 neurons in 2 birds, example raster Fig. 3a), higher than expected for an inhomogeneous Poisson process with identical rate modulations (Fig. 3b, mean recorded burst fraction = 0.19 ± 0.03, mean ± s.d., factor of 4.5 larger than expected, n=13 neurons in 2 birds; t_24_=15.0, p=6×10^−14^, one-sided two-sample t-test; Extended Data Fig. 4a-c).

**Figure 3:**
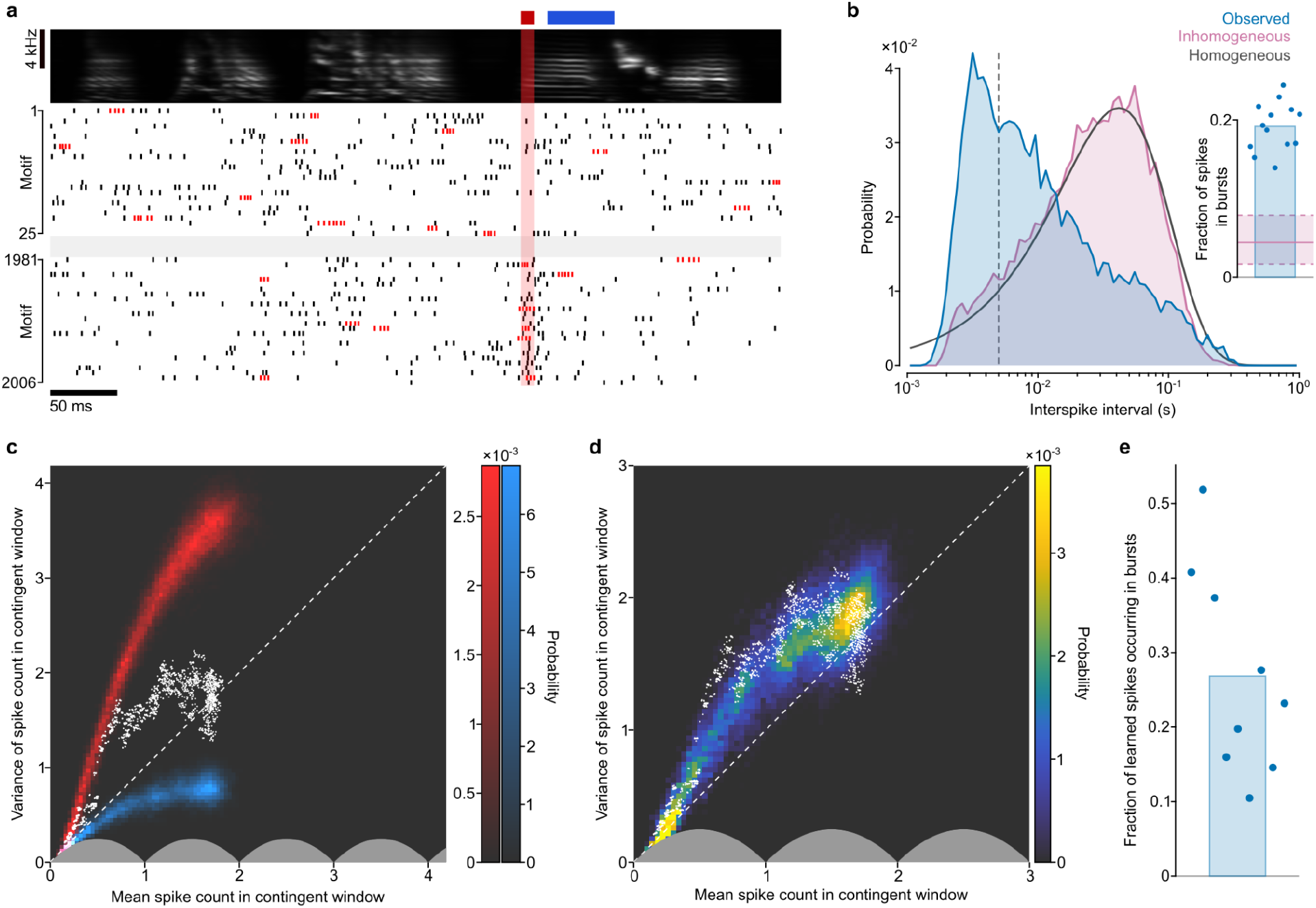
Learning is driven in part by an increased probability of burst events. **a**, Raster of a targeted single neuron for a set of motifs at the beginning (top) and end (bottom) of a day of nCAF learning (contingent window shaded in red; time of DAF marked in blue; detected bursts plotted in red, see Methods). **b**, Interspike interval distribution (excluding the contingent window) for the neuron in panel a (blue), a simulated inhomogeneous Poisson process with the same rate parameter modulation (pink), and a simulated homogenous Poisson process with the same average rate parameter (gray). Inset: fraction of spikes within burst events for a set of n=13 single LMAN neurons (blue; 2 birds). Pink shading shows the fraction of simulated spikes within bursts for a set of Poisson processes with the same rate parameter modulations (solid line is the mean across simulations for all neurons, dashed lines are 95% CI). **c**, White points: mean and variance of the spike count in the contingent window for the neuron in panel a, computed across a sliding block of 100 consecutive motifs. Heatmap: distributions of spike statistics in the contingent window for simulated models of learning using a pure bursting (red) or a pure inhomogeneous Poisson model (blue, see Methods). **d**, Same points as in panel c, plotted against a simulated spiking model of best fit that incorporates both bursting and Poisson firing. **e**, Contribution of bursting to nCAF learning for n=9 single neurons, computed using the composite model in panel d; plot shows fraction of learned spikes occurring within bursts (3 birds; see Methods).

To determine whether there was an increase in the probability of burst events in the contingent window, we considered spike statistics in the contingent window for a set of well-isolated, targeted nCAF neurons. We found that the variability of the spike count in the contingent window is too high to be explained by a Poisson model and too low to be driven entirely by bursts (Fig. 3c). It is instead consistent with a composite model (Fig. 3d; additional examples in Extended Data Fig. 4d-f), in which 27 ± 14% (mean ± s.d.) of increased spiking is due to an increase in bursting (Fig. 3e, n=9 neurons in 3 birds). This analysis suggests that, while bursts do increase slightly more than expected in the contingent window, they do not play an especially privileged role in nCAF learning.

### Learning is specific to single neurons

We next inquired about the spatial precision of nCAF learning—that is, to what extent are learned changes restricted to the targeted neuron? In our model, learned changes to the activity of an LMAN neuron are driven by a connection to this LMAN neuron from Area X, via DLM^32^. Consequently, the spatial precision of learning is limited by the precision of this feedback projection. Note that there is conflicting evidence as to whether the feedback projection from DLM to LMAN conveys only learned biases^40,41^ or also is a source of motif-to-motif variability in LMAN activity^42,43^. Here, we remain agnostic to the question of how LMAN variability arises and instead focus solely on how learned changes are distributed among LMAN neurons. At one extreme, each MSN in Area X could measure the activity of one LMAN neuron and transmit a learned bias only to that same LMAN neuron; in this case, nCAF learning might only occur in the targeted neuron (Fig. 4a). Alternatively, the feedback signal could be spatially diffuse, thus driving learning in an entire population of nearby, untargeted neurons (Fig. 4b). Prior axon tracing studies have placed an upper bound on the spatial precision of this feedback projection at approximately 200 μm^27^, but the exact value is not known.

**Figure 4:**
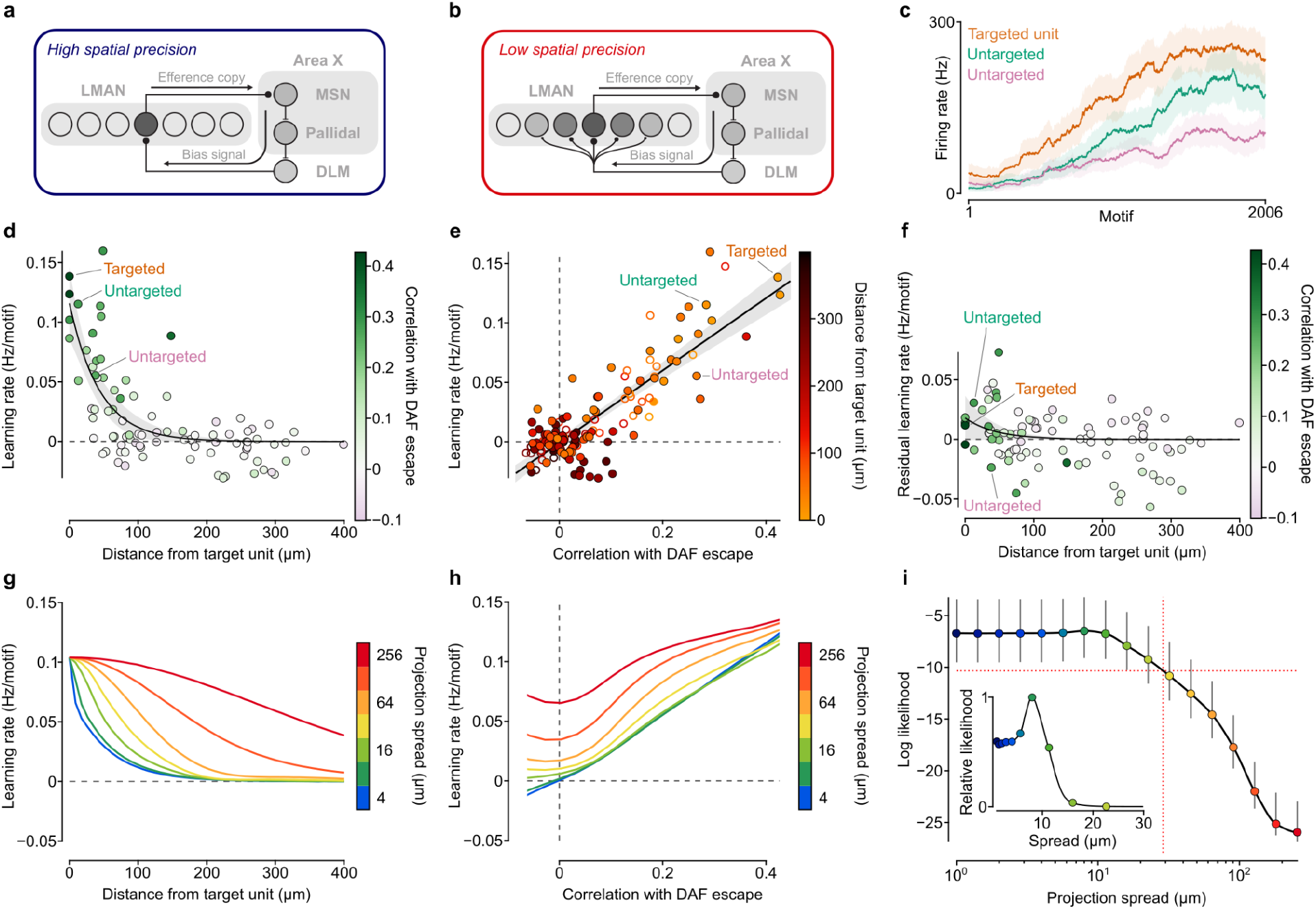
nCAF learning shows single-neuron specificity. **a**, With maximal precision, feedback learning projections could target single LMAN neurons, such that learning occurs only in those neurons correlated with DAF escape. **b**, Alternatively, feedback projections could spread widely over nearby neurons, such that learning is diffuse and non-specific. **c**, Firing rates over the course of an nCAF experiment for one targeted unit and two nearby untargeted units (firing rates calculated over a 100 motif sliding block; shading is 95% CI). **d**, Learning rates in many simultaneously recorded LMAN units (n=89 single units, 3 birds), as a function of their distance from the target neuron. Color represents correlation with DAF escape. An exponential fit is plotted in black (*λ* = 44 ± 3 um; shading is 95% simultaneous confidence band). **e**, Learning rates as a function of the correlation of each unit with DAF escape; color represents distance from the target neuron (3 birds, filled circles are single units, n=89; hollow circles are multi-unit signals, n=52; black line is linear fit: *β* = 0.30 ± 0.03 Hz/motif, 95% CI). **f**, Same single units as in panel d, but after subtracting the learning rate predicted by each neuron’s correlation with DAF escape, based on panel e linear fit. This result suggests that learning in neighboring neurons is largely explained by correlations with DAF escape. **g**,**h**, Model results for learning rates as a function of feedback projection spread (see Methods), **g**, Model learning rate vs distance from the simulated target neuron. Each curve represents the mean learning rate across all simulation results at a given distance from the target neuron. **h**, Same as panel g but with learning rates plotted as a function of correlation with DAF escape. Note strong correspondence between data in panels d,e and model results in panels g,h for small values of spread in feedback projections. **i**, Relative log likelihood that the observed single-unit data (same as in panel d) are generated by simulated models with a range of projection spread (Inset: zoom-in with linear scale; colors show correspondence with spread in panels g,h; red horizontal and vertical dashed lines indicate 95% confidence interval threshold).

We sought to distinguish between these two possibilities by quantifying learned changes in the activity of nearby LMAN neurons not targeted in our nCAF experiments. We observed learning in many nearby neurons (Fig. 4c,d). An analysis of the learning rates of LMAN neurons, as a function of their distance from the targeted neuron, reveals a spread of learning around the targeted neuron of 44 ± 3 μm (width to e^−1^ of exponential fit, 95% CI, n=89 single units in 3 birds, Fig. 4d), which is substantially smaller than the upper bound from anatomical studies.

This spread does not necessarily represent the fundamental spatial precision of learning in the AFP, however. Note that a substantial fraction of LMAN neuron pairs are significantly correlated within a range of around 100 μm (Extended Data Fig. 5a). As a result, some untargeted LMAN neurons are correlated with the targeted neuron, and thus correlated with DAF escape (Extended Data Fig. 5b-e). Thus, we expect that even if feedback projections to LMAN neurons had single-neuron specificity, these nearby correlated neurons should also learn. Indeed, across all neurons in our dataset, learning rates were positively associated with the degree of correlation between spike count in the contingent window and DAF escape, even in neurons that were not explicitly targeted with nCAF (Fig. 4e, r^2^=0.72, n=141 units in 3 birds). Finally, the degree of correlation with DAF escape predicts well the spread of learning in surrounding non-targeted neurons. After regressing out the effect of correlation with DAF escape, the amplitude of learning rates at short distances from the targeted neuron is reduced by a factor of 6 (scale factor of 0.16 ± 0.19 s.d., Fig. 4f). This result suggests that the degree of correlation with DAF escape is the primary determinant of learning rate, not proximity to the targeted neuron.

To more precisely determine the extent of the residual spatial spread, we fit our data to a quantitative model of learning that incorporates different degrees of spread in feedback projections (Fig. 4g-h, with a Gaussian distribution, see Methods). We found that our data are best explained by a model with a spread of 8 μm (0 to 29 μm 95% CI, n=89 single neurons in 3 birds, Fig. 4i). These findings are consistent with a neural circuit that precisely coordinates learned changes in activity with the correlation between neural activity and DAF escape. Nearby, uncorrelated neurons are excluded from incorrect learning with single-neuron precision.

### Learning is sensitive to feedback latency

We next inquired how the rate of nCAF learning depends on the time interval between the contingent window and the disruptive auditory feedback. One might imagine measuring this relationship by systematically shifting the interval between a narrow contingent window (*e*.*g*., 10 ms) and DAF; however, this approach would require many nCAF experiments over multiple days, during which drift in recording quality could confound the results. To overcome this challenge, we made the measurement in a different way; namely, we used a long contingent window (up to 120 ms) and made DAF playback contingent on the total number of spikes of an LMAN neuron across the entire window (Fig. 5a). As expected, we observed that neural activity was correlated with DAF escape across the entire contingent window (Fig. 5b). Thus, each timepoint in the contingent window is analogous to an nCAF experiment with a different DAF latency.

**Figure 5:**
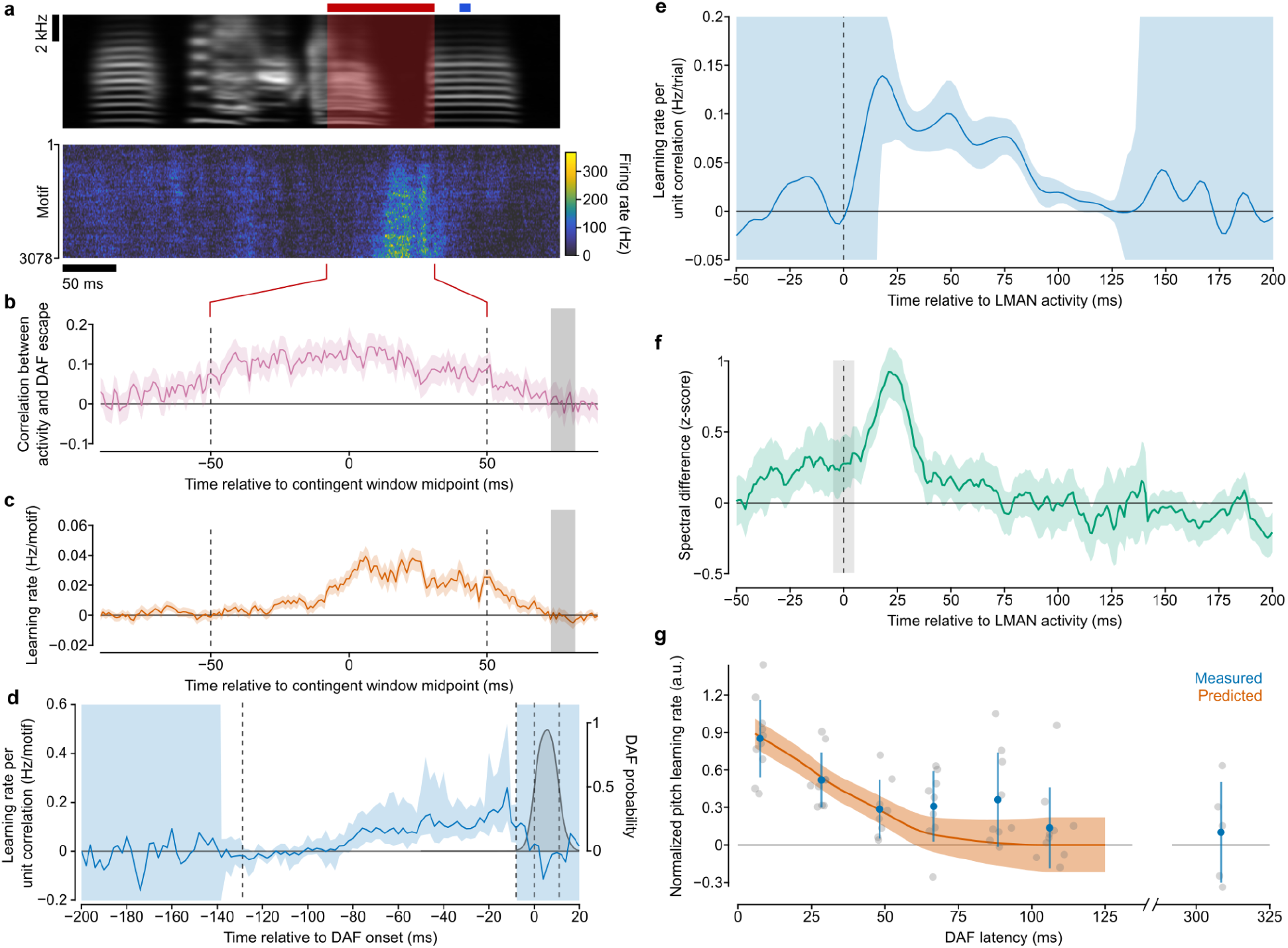
Direct measurement of eligibility trace within the vocal learning circuit. **a**, Results of an nCAF experiment where a short 10 ms DAF burst was played contingent on low activity in a long 100 ms contingent window. (Top panel) Song motif (contingent window shaded red; time of DAF in blue). (Bottom panel) Song-aligned firing rates of the targeted unit across a day of nCAF learning. **b**, Correlation between targeted neuron activity and DAF escape. **c**, Learning rate for a range of timepoints relative to the contingent window (vertical dashed lines; shading is 95% CI; DAF location plotted in gray). **d**, Learning rate per unit correlation, computed as learning rate from panel b, normalized by correlation from panel c, as a function of time in song (n=49 units in 3 birds; shading is 95% CI; see Methods). Gray shaded curve indicates DAF profile, i.e. the probability across motifs that a given timepoint overlaps with DAF. **e**, Inferred algorithmic eligibility trace (see Methods; shading is 95% CI). **f**, Measurement of premotor latency from LMAN activity to vocal output (n=3 birds). Green curve shows the magnitude of the effect on song of LMAN activity in a 10 ms window (gray) for different latencies (see Methods, shading is 95% CI). **g**, Normalized pitch learning rates for pCAF experiments at different DAF delays (n=60 experiments in 4 birds). Gray points are individual experiments. Blue points are means across experiments with similar delays. Error bars are standard deviations. Orange curve is the predicted pCAF learning computed from the measured LMAN latency (panel f) and the measured algorithmic eligibility trace (from panel e) (see Methods; shading is 95% CI).

Despite the correlation between DAF and neural activity extending over the entire contingent window, we found that learning was restricted to a much narrower range of times, late in the window (Fig. 5c; Extended Data Fig. 6). From these data, we extracted the learning rate per unit correlation (*normalized* learning rate) for each time in the contingent window (Fig. 5d). Using this normalized learning rate, we were then able to infer a function that describes the eligibility of LMAN neurons to learn from acoustic feedback at different delays. We refer to this function as the *algorithmic eligibility trace* (see Supplementary Note) because it describes the relationship between LMAN learning and the time of the auditory feedback, without reference to a specific biological implementation (*e*.*g*., a synaptic eligibility trace and dopamine signals). We estimate this algorithmic eligibility trace by modeling our normalized learning rate curve as the convolution of the algorithmic eligibility trace with the average DAF timecourse. Using deconvolution, we obtained an algorithmic eligibility trace that is significantly nonzero only up to a delay of 126 ms after LMAN activity (95% CI = 108-140 ms, n=49 units in 3 birds, Fig. 5e).

One might expect that the algorithmic eligibility trace would overlap with the vocal premotor delay of LMAN neurons, so that the song system can learn from song variations generated by LMAN. To test this idea, we computed the magnitude of differences in the song spectrogram corresponding to changes in LMAN activity (Fig. 5f, Extended Data Fig. 7, see Methods). We found that the greatest difference in song occurs at a latency of 22 ms (95% CI 20-24 ms) relative to LMAN activity. This is well aligned with the maximum measured value of the algorithmic eligibility trace. Notably, this premotor latency is shorter than what has been previously estimated using microstimulation experiments (35-70 ms^44^).

Finally, we wondered if we could use the estimated algorithmic eligibility trace and premotor latency to predict the dependence of pitch-conditional auditory feedback (pCAF) learning on the timing of DAF playback. We performed a series of pCAF experiments with different DAF delays and found that the measured pCAF learning rates are consistent with those predicted from our nCAF experiments (Fig. 5g, 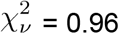, n=60 experiments in 4 birds, see Methods) and with past experiments^29^.

## Discussion

Here we demonstrate that the song learning circuitry biases LMAN activity towards patterns associated with improved vocal performance, confirming a key untested prediction of our circuit model. We further show that this credit assignment operates at the level of individual LMAN neurons with millisecond-scale temporal precision. This degree of spatiotemporal specificity strongly constrains the possible circuit wiring and biophysical mechanisms that underlie credit assignment in the songbird cortico-basal ganglia circuitry.

In our circuit model, medium spiny neurons (MSNs) in Area X receive three distinct inputs: timing information from HVC; an efference copy of LMAN activity; and a report of relative song quality from VTA in the form of a diffuse dopamine signal (Fig. 6a). The first component of our model is that coincident HVC and LMAN inputs onto an MSN establish an eligibility trace at the HVC-to-MSN synapse. In one possible implementation of this coincidence detection^45^, HVC projections synapse onto NMDA-receptor-predominant spines and LMAN inputs fall onto AMPA-receptor-predominant dendritic shafts. When an HVC neuron bursts, glutamate binds to NMDA receptors on the spines; however, voltage-dependent magnesium block prevents current flow through these channels. Only when the LMAN neuron fires—depolarizing the dendritic segment and thereby relieving this block—does calcium enter the spine. This calcium influx could tag the spine with a synaptic eligibility trace, which acts as a transient record that the LMAN neuron was active at the time encoded by this HVC input (Fig. 6b). The second component of the model is that a subsequent increase in dopamine from VTA, signaling better than average song performance, strengthens the tagged HVC-to-MSN synapse (Fig. 6c). In this way, LMAN activity that improves the song at a particular time leads to strengthening of the synapses from the HVC neurons that encode that time onto the MSNs associated with that LMAN neuron. On later song renditions, the strengthened HVC input drives the MSN to fire at this same timepoint, reactivating the same LMAN neuron via closed topographic loops through the Area X→DLM→LMAN pathway and thereby reinforcing the activity associated with improved performance. Notably, this circuit model provides a mechanism for credit assignment with high spatiotemporal precision, despite diffuse, delayed dopamine signals^32,46^.

**Figure 6:**
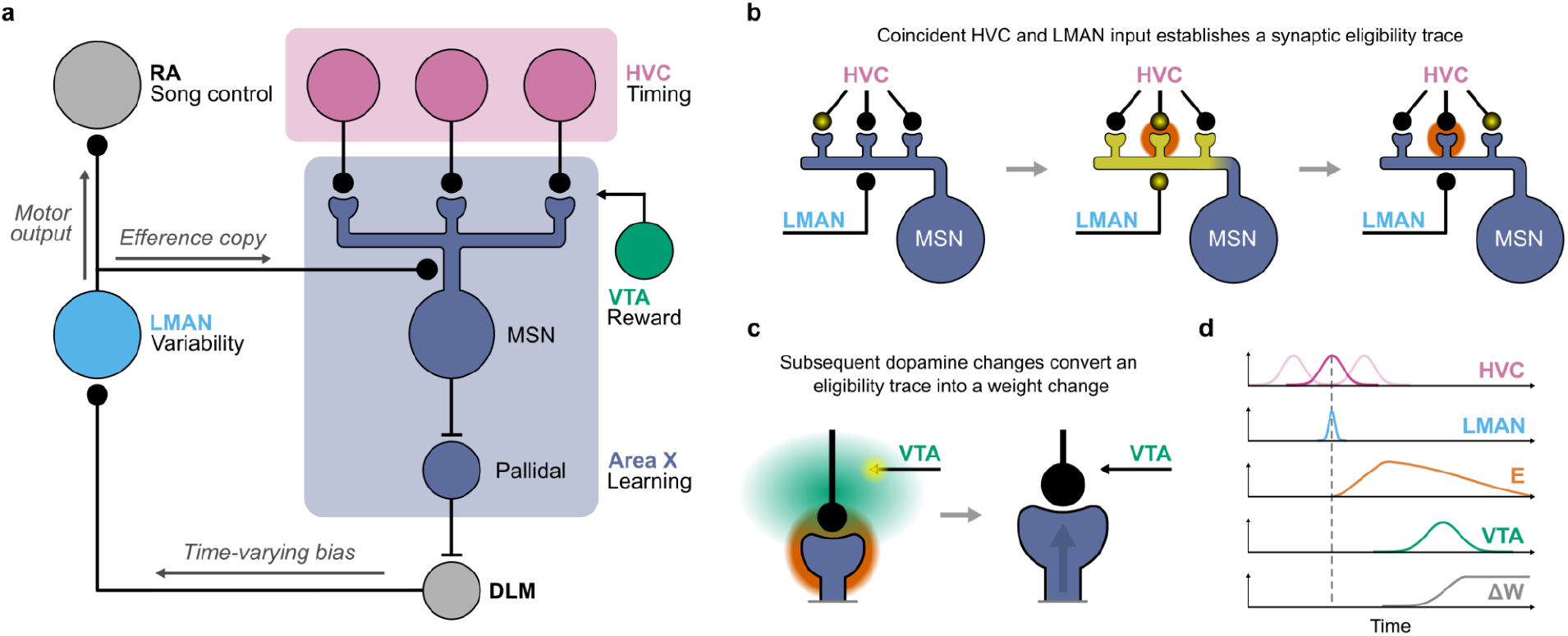
A circuit model for basal-ganglia-mediated reinforcement learning in the songbird. **a**, Circuit diagram of the vocal learning circuitry. Depicted are: temporal sequence generator HVC; primary motor cortex analog RA; variability-generating nucleus LMAN; basal ganglia nucleus Area X, containing a medium spiny neuron (MSN) and pallidal neuron; thalamic nucleus DLM; and the ventral tegmental area (VTA). **b**, Model of synaptic eligibility trace formation within Area X MSN spines^45^. HVC neurons, which synapse onto dendritic spines, fire sequentially during singing. Spiking of an LMAN neuron, which synapses onto the dendritic shaft, depolarizes the entire dendritic segment and establishes a transient synaptic eligibility trace within the spine(s) that received co-active HVC input (represented as orange glow). **c**, HVC-to-MSN synapses tagged with an eligibility trace strengthen in response to subsequent dopaminergic input from VTA. On subsequent song renditions, these strengthened synapses cause the MSN to fire at the time encoded by the strengthened HVC input, reactivating the LMAN neuron that was associated with improved song performance at that time. **d**, Timing diagram of the proposed three-factor, corticostriatal plasticity rule at Area X MSNs. Our findings suggest that the high temporal resolution of learning is supported by a precise coincidence detection between HVC and LMAN inputs. ‘‘E” represents the synaptic eligibility trace and “ΔW” represents the change in weight of the HVC-to-MSN synapse.

The millisecond-scale precision of nCAF learning demonstrates that the song learning circuitry faithfully resolves the timing of individual LMAN spikes within song. Notably, the width of the learning kernel (7.5 ms full width half maximum = 3.2 ms s.d.) is comparable to the timescale of an HVC_X_ neuron burst (6.3 ms width, ± 1 ms onset jitter^47^), indicating that temporal resolution is preserved with high fidelity through the coincidence detection step. This is consistent with—but near the limit of—NMDA receptor-mediated coincidence detection, which has been shown to act on timescales of ~5-30 ms^45,48–50^. Thus, if coincidence detection by an Area X MSN utilizes NMDA receptors, it may require unique specializations.

In addition to detecting the coincidence of HVC and LMAN spiking with low temporal jitter, the AFP needs to reactivate the LMAN neuron at the correct time. Our measurements suggest that the AFP can achieve this to within an offset of 3.0 ± 2.3 ms (mean ± s.d.). This requires the AFP to overcome an additional challenge—the propagation delay between Area X and LMAN. That is, to drive LMAN spiking at a desired time in the song, an MSN must fire *before* this time by an amount that compensates for the travel time through the multi-synaptic pathway between Area X and LMAN. To achieve this, the HVC-LMAN coincidence detector could respond preferentially to HVC inputs followed by LMAN inputs with a delay equal to the total propagation time from LMAN to the MSN, through the AFP, and back to LMAN, reminiscent of plasticity in mammalian cerebellum^51^.

We also find that nCAF learning is targeted to the correct LMAN neurons with a spatial precision on the order of a single cell; however, care must be taken in interpreting this result. Note that in nCAF, correlations between nearby LMAN neurons mean that many neurons are correlated with the targeted neuron and with DAF escape, and thus should learn. With this in mind, our data provide strong evidence against nonspecific, divergent projections in the AFP that are shared across correlated and uncorrelated LMAN neurons and support a more precise pattern of projections. However, we cannot determine whether an ensemble of correlated LMAN neurons receives a shared feedback projection, or if each neuron receives an independent projection from DLM. In fact, if LMAN variability is driven primarily by variable input from DLM, then we would expect LMAN neurons that share a projection to be correlated with each other. This uncertainty could potentially be resolved by a connectomic reconstruction of LMAN (including incoming DLM axons) after functional imaging. In either case, it is an intriguing question how this wiring specificity is achieved, whether through molecular guidance^52^, spontaneous activity^53^, or experience^54,55^.

In this study, we have introduced the concept of an algorithmic eligibility trace, which describes the eligibility of a neuron to learn in response to errors or rewards at different latencies, without reference to a specific circuit implementation. In our model, however, we expect that the algorithmic eligibility trace is implemented by a synaptic eligibility trace in MSN spines that responds to subsequent dopamine signals from VTA. The algorithmic eligibility trace allows us to make predictions about this synaptic eligibility trace (Supplementary Note). In particular, we found that the offset of the algorithmic eligibility trace—the longest delay of auditory feedback that can drive learning—is 126 ms after LMAN activity. Previous measurements have shown that VTA neuron response onsets to acoustic noise bursts are delayed by ~55 ms^8^ (relative to noise onset). Thus, if LMAN neurons can learn from auditory feedback delayed by up to 126 ms, and the dopaminergic signal corresponding to such auditory feedback is delayed by an additional 55 ms, then the offset of the synaptic eligibility trace should not occur until 181 ms after LMAN activity. This is much shorter than the seconds-long eligibility traces measured at mammalian corticostriatal synapses^13,56^. Unfortunately, we are unable to observe the onset of the algorithmic eligibility in our experiments because we cannot play DAF contingent on LMAN activity prior to the occurrence of that activity. Therefore, we cannot infer the time of onset of the synaptic eligibility trace. This question can be resolved by future experiments where dopamine levels in Area X are directly manipulated (*e*.*g*., using optogenetics) contingent on LMAN activity.

The nCAF paradigm extends a fruitful tradition of leveraging brain-computer interfaces to probe mechanisms of learning in the brain^57–59^. What distinguishes nCAF from prior closed-loop paradigms is the unique interpretability provided by the song learning system. Reinforcement learning is fundamentally a process of mapping states to actions; however, in many BCI paradigms the internal state representation being used by the learning circuitry is unknown. By contrast, there is an abundance of evidence that in song learning, the relevant state is time in song, as encoded by HVC^20,21,60^. It is due to this known state representation, along with the constrained circuit wiring of the song system, that our observations provide interpretable constraints on the song learning circuitry.

Together, these results establish that the avian cortico-basal ganglia circuit for vocal learning solves the credit assignment problem with a remarkable degree of spatiotemporal precision. Learning is targeted to the correct premotor neuron with single-neuron specificity, and to the correct time in song with millisecond-scale precision. This specificity is consistent with our circuit model for basal-ganglia-mediated learning and suggests that each aspect of the circuit is highly tuned to meet the demands of song learning. Whether mammalian cortico-basal ganglia circuits are able to learn with similar precision remains an open and important question.

## Methods

### Animals

Male zebra finches (*Taeniopygia guttata*) were obtained from the Massachusetts Institute of Technology zebra finch breeding facility (Cambridge, MA). Animals were housed under a 12h/12h day/night cycle at 24°C and 35% relative humidity. Animal care and experiments were carried out in accordance with the guidelines of the National Institutes of Health and were reviewed and approved by the MIT Committee on Animal Care (protocols 0718-066-21 and 2404-000-657). Birds were raised with both mother and father present until approximately 55 dph to allow for naturalistic tutoring. Young adult male zebra finches (>90 dph) were separated from the colony and singly housed in acoustic isolation chambers. Those birds who showed the greatest proclivity to sing were selected for experimental use.

### Surgery

Birds were given a preoperative intramuscular injection of meloxicam (1-2 mg/kg) and anaesthetized with 1-3% isoflurane gas before being placed in a custom stereotaxic apparatus. Feathers were removed from the scalp and the skin disinfected using povidone-iodine. A subcutaneous bupivacaine injection (3 mg/kg) was administered to the scalp and a midline incision was made using surgical scissors. Craniotomies were made using a high-speed rotary dental drill (Dentsply Sirona) at locations measured relative to the lambda sinus, a major blood vessel. Craniotomies were approximately 1000 um AP x 400 um ML. Neuropixels implants were inserted using known anatomical coordinates for LMAN (head angle 20°, AP 5.3 mm, ML 1.8 mm, DV 1.9 mm). Head angle was measured relative to a horizontal defined by the flat skull surface immediately posterior to the beak. AP and ML coordinates were defined relative to an origin set at the lambda sinus, measured through the inner leaflet at a 20° head angle. DV coordinates were defined relative to the surface of the intact dura measured immediately after craniotomy. Microdrives were inserted with the Neuropixels shanks in the retracted position. The origin of the implant was set at the midpoint between the tips of the two central Neuropixels shanks. Shanks were inserted at a rate of 100 μm/min using a custom motorized stereotaxic drive to a depth of 1.0 mm DV. Prior to insertion, shanks were coated in DiI (1,1’-dioctadecyl-3,3,3’,3’-tetramethylindocarbocyanine perchlorate, Invitrogen) for later histological identification. Craniotomies were sealed using Dura-Gel (Cambridge Neurotech) and implants were anchored to the skull using a combination of dental acrylic and C&B Metabond. The incision site was then closed using veterinary cyanoacrylate adhesive (3M Vetbond) and anaesthesia was removed. Birds were administered postoperative meloxicam twice a day for two days following surgery. Once birds had fully recovered and resumed singing at their pre-surgery rates, the Neuropixels shanks in the microdrive were advanced by 2.5-3.0 mm, so that they passed through LMAN.

### Histology

In a subset of implanted birds, we verified the placement of the Neuropixels with histology. Following transcardial perfusion with paraformaldehyde (Sigma), sections were cut on a vibrating microtome (Vibratome), stained with NeuN (conjugated to AlexaFluor 647), and visualized using an Axioplan 2 fluorescence microscope (ZEISS). In all birds examined, the location of LMAN aligned with that found using electrophysiological features (see below).

### Data acquisition

Recordings were performed using SpikeGLX software (https://billkarsh.github.io/SpikeGLX) and National Instruments PXIe hardware (PXIe-1071 or PXIe-1083 chassis). Electrophysiology data were acquired using Neuropixels 2.0 probes^61^ in a custom microdrive^35^. Probes were connected to an assisted rotary joint (Doric AERJ_24_HDMI+4) in a sound isolation chamber to allow for free movement of implanted birds, and then connected to a Neuropixels Basestation (IMEC PXIE_1000). Audio was recorded using an electret microphone (GRAS 40AE), amplified with a Focusrite Scarlett 2i2 USB audio interface and fed into a National Instruments analog-to-digital converter (PXIe-6341 with a BNC-2110 breakout).

### Neuron-conditional auditory feedback (nCAF)

Closed-loop control of data saving and disruptive auditory feedback (DAF) delivery was achieved using custom workflows written in Bonsai^62^.

#### Song-triggered saving

In a subset of recordings, audio and Neuropixels data were only saved during periods when the bird was singing. Audio from SpikeGLX was continuously streamed into the Bonsai workflow using the Bonsai.SpikeGLX package. Song bouts were detected by monitoring sound power in the band from 500 to 8000 Hz. Singing was defined as a period when the average power in this band exceeded a user-defined threshold. When a song bout was detected, a digital line was set high on a National Instruments USB device (USB-6008 or USB-6501). This line was used as a TTL trigger by SpikeGLX to initiate saving of a new file. Importantly, the saved data included 3 seconds prior to the digital line being set high and 3 seconds after it being set low.

#### Contingent time detection

Disruptive auditory feedback was targeted to a specific time in song by detecting the combination of acoustic features associated with this time. The song-feature detector consisted of two components: (1) an initial template matching detector that recognized a sequence of syllables from their associated spectrogram, and (2) a syllable onset detector that was armed when a template match was detected and triggered on the rising edge of the next syllable. This two-part system allowed for more precise detection of time in song than a template-matching approach alone.

#### Online spike inference

Once the contingent time in a song motif was detected, a decision was made about whether to play DAF. This was achieved as follows. First, a brief snippet of electrophysiology data, was fetched from SpikeGLX. Typically, this snippet was the most recent *L* ms of data from SpikeGLX at the moment the contingent time was detected, where *L* ms was the length of the contingent window. For nCAF experiments with a 5 ms contingent window (Fig. 2), targeting jitter was further reduced as follows. Shortly after the contingent time in song was detected using the method described above, long (~1 second) snippets of both audio and Neuropixels data were fetched from SpikeGLX. These snippets were aligned using the 1 Hz sync wave recorded on both the Neuropixels device and the NI-DAQ. The onset time of the targeted syllable was found in the audio snippet and then used to locate the contingent window in the Neuropixels snippet.

The electrophysiology data snippet was then processed to estimate the number of spikes from the targeted neuron within the contingent window. First, the snippet underwent common-average referencing (CAR) to remove common-mode noise and artifacts. The snippet was then filtered using a 2nd order high-pass Butterworth filter with a cut-off frequency of 300 Hz to isolate the action potential frequency band. Note that the snippet was padded with reflections on both ends to minimize edge effects, and the filtering was performed in both a forward and backward pass to prevent introducing phase offsets. In early experiments, we observed that when the snippets contained many spikes—which are each dominated by a large negative deviation—this high pass filtering introduced a positive DC offset relative to the “true” voltage baseline. To remedy this, we subtracted the median value of each filtered snippet.

After this preprocessing was complete, the number of spikes the targeted neuron generated in the contingent window was estimated in one of two ways: simple threshold crossing, or template matching.

##### Threshold crossing

In the case of threshold crossing, the number of spikes was estimated as the number of times the signal crossed a predetermined threshold, *T*. This threshold was computed using the method in ^63^. Briefly, a calibration time-series of filtered data from the targeted channel, *x*, was acquired at the beginning of the experiment. The spike threshold was then defined as follows:

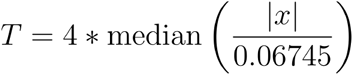

##### Bayes optimal template matching (BOTM)

The template matching method identified spikes by convolving voltage data with a precomputed filter, optimized to extract the activity of a specific neuron, following a Bayes Optimal Template Matching (BOTM) technique^64^. Briefly, an initial calibration dataset was acquired and spike sorted using the pipeline described below at the beginning of the day. Sorted units were manually curated and one was selected as the target neuron. Optimal filters and threshold for detection of this unit were generated using a custom MATLAB script and then supplied to the Bonsai workflow.

#### Disruptive auditory feedback (DAF)

Generation of acoustic white noise bursts was achieved using an Arduino UNO Rev3 flashed with a custom sketch. Briefly, when the contingent time in song was detected by Bonsai, a digital line was set high using a National Instruments USB DAQ (USB-6008 or USB-6501). This rising edge was detected by the Arduino and initiated a fixed-duration timer. If this timer elapsed without additional input, no noise was played. If a second digital line was set high during the timer interval (indicating a decision from the Bonsai workflow to play DAF), a fixed-duration period of white noise was generated once the timer elapsed. White noise was generated by repeatedly, pseudo-randomly setting the state of a digital output line on the Arduino using a 16-bit Galois linear-feedback shift register. The output of this digital line was amplified by a Pyle PFA300 audio amplifier and played through a Pyle PDMR5 speaker. The amplifier gain was manually calibrated using an EXTECH sound level meter to ensure that the acoustic noise bursts remained between 90 and 110 dB SPL.

### Pitch-conditional auditory feedback (pCAF)

Closed-loop control of data saving and disruptive auditory feedback (DAF) was achieved using custom workflows written in Bonsai^62^. Song-triggered saving, contingent time detection, and DAF generation were performed as in nCAF. Pitches were computed from a 10 ms snippet of audio by identifying the peak of the audio cepstrum within a user-defined quefrency range. For each bird, pCAF was performed over multiple days, with different DAF delays. The contingency direction (driving pitch up vs. driving pitch down) was swapped each day. Experiments with the same DAF delay were generally performed on consecutive days.

### Data analysis

#### Offline spike sorting

High density electrophysiology data were sorted using a custom analysis pipeline constructed in SpikeInterface^65^ to emulate the approach used by the International Brain Laboratory (IBL)^66^. First, CatGT (https://billkarsh.github.io/SpikeGLX/#catgt) was used to concatenate individual files, correct for phase shifts introduced by asynchronous analog-to-digital conversion of different channels, and bandpass filter traces between 300 and 9000 Hz using a 12th order Butterworth filter. Using SpikeInterface, bad channels were detected and interpolated, and a 3rd order spatial high-pass Butterworth filter (cutoff at 0.01 Nyquist) was applied. Motion correction was then performed using DREDge^67^. Spike sorting was performed using Kilosort 4^68^. Sorted units were manually curated in Phy (https://github.com/cortex-lab/phy). For analyses that required well-isolated single units (Fig. 1 where specified, Fig. 3 & Fig. 4d,f,i) the median spike amplitude, amplitude cutoff, and sliding refractory period violation metrics used by the IBL^66^ were used to identify putative single units. Units that passed these criteria were then manually inspected to verify appropriate waveform shape, refractory periods, and stability over the recording duration. For analyses that were less sensitive to unit isolation (Fig. 1 where specified, Fig. 2, Fig. 4e & Fig. 5), we used units with a median amplitude greater than 50 mV that passed a visual inspection for waveform shape, without regard to refractory period. The targeted unit in each experiment was identified from the spike sorting output by finding the unit with the greatest waveform similarity to the filter used for online spike detection (in the case of template matching) or the largest spike amplitude on the targeted channel (in the case of threshold crossing).

#### Acoustic analysis

Spectrograms were calculated using the short-time Fourier transform with a Hamming window function and a window width of 10 ms (using the MATLAB spectrogram function). Songs were aligned relative to the onset of the syllable containing the contingent window. For analyses that were sensitive to variations in song element length (Fig. 1c-g, Fig. 5f, Extended Data Fig. 7), songs were time warped and aligned to each syllable onset and offset^6^. All spectrograms are plotted on a logarithmic scale.

#### LMAN localization

Identification of the boundaries of LMAN in our recordings was achieved using a two step method. First, approximate LMAN boundaries were determined using a measure of song modulation of multi-unit spiking activity on each channel as follows. Multi-unit activity on each pad was estimated using the same threshold crossing technique described in Methods section Online Spike Detection, and aligned to song. A time-dependent multi-unit firing rate was then computed by smoothing spike trains with a 2 ms Gaussian kernel and averaged across song motifs to yield a song-locked firing rate *r*_*i*_(*t*) for each channel, *i*. A song modulation metric was then calculated as

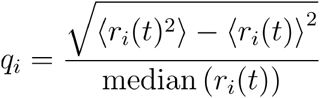

A threshold value of *q* was calculated using Otsu’s method^69^, and the largest contiguous region of pads with values of *q*_*i*_ larger than this threshold was used as an initial estimate of LMAN.

The initial estimate of the boundaries of LMAN was refined using a classifier based on the waveforms of each sorted unit on the Neuropixels shank. The initial LMAN boundaries were reduced by 25% to define a region of units definitely within LMAN. Similarly, the initial bounds were expanded by 25% and the region beyond this bound was defined as definitely not containing LMAN units. After using the synthetic minority oversampling technique (SMOTE) to balance classes, the waveform features of units in each of these regions (waveform width, positive peak, and negative peak) were used to train an SVM. This SVM was then used to compute class scores for all units on the shank, which were converted to posterior probabilities using a sigmoid function. An exhaustive search was then performed over all possible boundary locations to find the maximum likelihood LMAN boundaries. (Fig. 1, Fig. 5),

#### Learning rate estimation

Learning rates were computed by fitting a linear model to firing rate as a function of motif number. For measurements of learning profiles (Fig. 1g, Fig. 2, Fig. 5), this fit was performed independently at each time point after downsampling the raw spike times into 0.5 ms bins. For overall learning rates (Fig. 1d,f,h, Fig. 3, Fig. 4), a linear fit for each unit was computed using the total number of spikes within the measured time window on each motif. Confidence intervals were constructed using the standard error of the model slope.

#### Learning kernel measurement

Learning precision was estimated from learning profiles and correlation profiles. Learning rates were estimated using the linear fit procedure described above. Correlations were estimated by calculating the Pearson correlation coefficient across motifs between the spike count in each time bin and a Boolean variable representing if noise was played on each motif. Confidence intervals were calculated using a Fisher transformation.

Gaussian distributions were fit to both the correlation profile and learning profile curves to estimate the width (*μ*_*c*_ and *μ*_*r*_) and relative time offset (*σ*_*c*_ and *σ*_*r*_) of each. Confidence intervals on these parameters were calculated by bootstrapping (n=1000) with simulated correlation and learning profiles resampled using the uncertainties estimated for each curve.

For each unit, the position of the learning kernel *μ*_*k*_ was estimated as *μ*_*k*_ − *μ*_*k*_, and the population estimate as the mean of all values. The width of the learning kernel was estimated as 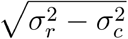. The population estimate was then calculated by fitting a hyperbolic function 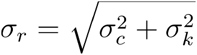 to the data (Fig. 2d), with confidence intervals calculated by bootstrapping (n=1000) with the uncertainties for each *σ*_*c*_ and *σ*_*r*_

#### Burst detection

We identified bursts in the spike trains of recorded LMAN units using the nonparametric approach described by Gourévitch and Eggermont^70^ with a rank-surprise threshold corresponding to a probability of 0.05 and an initial ISI threshold of 5 ms.

#### Bursting neuron simulation

We compared our observed spike statistics to simulated neural spike trains generated using both an inhomogeneous Poisson process and a variable bursting model. For some analyses, these two models were used independently (Fig. 3c). In another analysis, they were combined into a composite model of regular spiking and burst spiking (Fig. 3d,e).

In the first model we considered, changes in the firing rate of an LMAN neuron were driven entirely by rate modulations of a Poisson process. The Poisson process was simulated by estimating the time-dependent firing rate of a given unit within each song motif in a sliding window of 1 ms. The simulation also included changes in firing rate over the course of the day in a sliding window of 100 motif renditions. This time-dependent firing rate, which we denote *r*(*t, i*) where *t* is time in song and *i* is the song motif number, was then used to simulate a spike train. We used a refractory recovery function *w*(*t*) estimated from the data^71^. With this approach, we were able to simulate spike trains under different assumptions about the underlying statistics, as in Fig 3b. It was also used to simulate firing rates within the contingent window as in Fig 3c.

We also considered another model in which the learned increase in spike count in the contingent window was due to increased probability of bursting. In this model, we first estimated the background non-burst firing rate in the contingent window at the beginning of the day. We also estimated the firing frequency of the unit during burst events from the data (typically 300-500 Hz). In the simulated spike train, each motif was designated as either a burst motif (with probability) or a non-burst motif (with probability 1 − *p*). On burst motifs, spikes were generated using a Poisson process with the burst firing rate. On non-burst motifs, spikes were generated using a Poisson process with the background firing rate. The probability, *p*, was chosen so that the average firing rate of the model unit within the contingent window would match the average firing rate of the observed neuron over the course of learning. The resulting spike statistics of the bursting model (variance vs mean of spike count) were compared to the observed data.

The composite model explained increases in firing rate over the course of feedback as being driven by both Poisson and variable bursting contributions. Firing rate in the contingent window on a given motif was modeled as a baseline firing rate *F*_0_ plus a learned firing rate on the *j*th motif Δ*F*_*j*_. Both the Poisson firing rate *F*_*poisson*_ and the burst probability *p*_*burst*_ were scaled linearly with Δ*F*_*j*_, with the relative contribution controlled by a scalar parameter *β*. Thus,

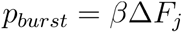

whereas Poisson firing rate scaled as

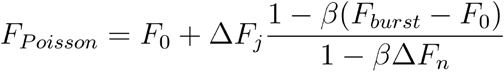

which results in the simulated firing rate equalling *F*_0_ + Δ*F*_*n*_, the observed firing rate at the end of feedback. The fraction of learned spikes in the contingent window generated by bursts is then given as

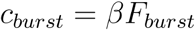

We fit *β* to each unit by maximizing the log-likelihood of our observed data. For a given nCAF experiment, we computed the mean and variance of the number of spikes in the contingent window across the experiment using a sliding window of 100 motifs. We then computed a probability distribution of predicted means and variances from simulated data. For a given *β*, we simulated a series of spikes that we used to compute a 2D probability distribution of the mean and variance of spike count. We could then vary *β* to maximize the likelihood of observing our data given this underlying probability distribution. Uncertainty in *β* was quantified by bootstrapping (n=100) using resampled mean and variance estimates.

#### Spatial precision simulation

We created simulated models of learning in LMAN neurons with different degrees of feedback spread, to see which of these models fits the data best. The model incorporated a realistic spatial distribution of LMAN neurons, and included pairwise correlations between LMAN neurons based on measured values. The learning rate of each model neuron was assumed to have a component due to its own correlation with DAF escape, and a component due to the spatial spread of learning from nearby, correlated neurons.

Locations of LMAN neurons and the pairwise correlations between them were extracted from the data as follows. The spatial location of each recorded unit was determined using monopolar triangulation with SpikeInterface. The targeted unit was identified by finding the unit with the largest spatial overlap with the filter used for online spike detection. The location of this unit was set as the origin, and the distances of other units from the targeted unit were calculated as the Euclidean distance from this origin. Pairwise correlations between units were calculated over a series of undirected motifs prior to the beginning of nCAF. Firing rates were calculated using a sliding window of 20 ms, and motif-by-motif firing rate fluctuations were estimated by subtracting a mean song-locked firing rate profile from the activity for each motif. The Pearson correlation coefficient was then calculated between the rate fluctuations for each pair of neurons across all times in song and motifs. A significance threshold was determined by shuffling the data in time using a Bonferroni correction for multiple comparisons.

Simulated learning rates were generated from a simulated population of LMAN neurons arranged on a grid with 20 μm spacing and 200×200×400 μm dimensions in x, y, and z. Grid locations were randomly jittered by 5 μm in each simulated population to allow for dense spatial sampling. We placed a simulated recording probe oriented along z at the x,y center of this volume, with the face of the probe in the yz plane. We then divided the population into two classes, recorded and unrecorded neurons. This division was done to account for the different statistics governing correlation with noise for recorded and unrecorded units, as a result of crosstalk in our feedback targeting. The recorded volume extended out from the active face of the probe by 20 μm in x and was 40 μm wide in y.

The simulated neuron at the center of the volume was designated the target neuron, and its correlation with noise was assigned by sampling from a Gaussian distribution of correlations based on recorded target neurons. The correlations of other neurons in the recorded volume were assigned by sampling from a Gaussian distribution with a mean and width that both decayed exponentially with distance from the target neuron. Both decay functions parameterizing this distribution were fit to the observed data (Extended Data Fig. 5d).

Correlations with noise for unrecorded neurons were drawn using a different process, since we had no ground truth for the correlation of unrecorded neurons with noise bursts. We instead first simulated their pairwise correlation with the target neuron, which we could model from our observed data. We could then take advantage of the fact that correlation with the target neuron scaled linearly with correlation with noise (Extended Data Fig. 5b) to convert this pairwise correlation into a correlation with DAF. Pairwise correlations were drawn from a distribution of correlation as a function of distance that was obtained from a kernel density estimate of the observed pairwise correlations. These pairwise correlations were then multiplied by a scalar factor determined by a fit to the data (Extended Data Fig. 5b) to yield a correlation with DAF for each simulated neuron.

Once correlations with DAF were assigned for each simulated neuron, we then calculated the predicted learning *r*_*i*_ of each neuron for a given value of *σ*_*d*_, the spatial divergence of projections in the circuit. We modeled this divergence as a Gaussian, yielding an expression for the learning for each neuron as

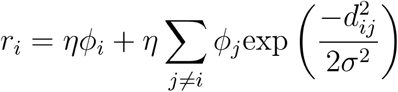

where *ϕ*_*j*_ is the correlation of each neuron with DAF, and *d*_*ij*_ is the distance between each pair of neurons. Note that this can we written more succinctly as

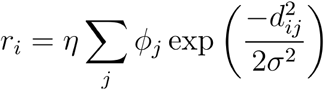

Since *d*_*ij*_ = 0 when *i* = *j*.

We then used our simulation results to construct, for a range of values of *σ*_*d*_, a 3D probability distribution of predicted learning rates at each combination of distance and correlation, using additive smoothing with a Jeffreys prior of *σ*= 0.5. We could then compare these distributions with our observed data to calculate the relative likelihood of our data having been generated by the distribution corresponding to each value of *σ*_*d*_. Each value of *σ*_*d*_ was simulated n=10^4^ times to generate the results in Fig. 4. Error bars were determined with a bootstrap (n=10^4^) using uncertainties in the estimates of correlation and learning rate in the data.

#### Eligibility trace estimation

By using a long contingent window before the DAF noise burst (Fig. 5a), we were able to induce a correlation between neural activity and DAF escape that extended over a 100 ms or longer period. This led to learning restricted to the second half of the contingent window, suggesting that correlations in the first half of the contingent window were too early before DAF, and fell outside a window for sensitivity of the learning circuit. The shape of this window of sensitivity is referred to as the algorithmic eligibility trace (see Supplementary Note). To extract this eligibility trace, we started by computing the normalized learning rate, as follows (Fig. 5e).

First, we note that, at short latencies before DAF, learning rate is roughly proportional to the correlation of the neural activity with DAF escape (Fig 4e). Thus, we normalized learning rate at each point in time by the correlation with DAF escape of activity at that time. This normalized learning rate, 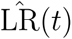, was calculated as the median of the set of ratios between learning rate and correlation across all units. Confidence intervals were obtained with a hierarchical bootstrap (n=1000 replicates), first resampling with replacement across birds and then jittering learning rates and correlations based on their uncertainties. This calculation was repeated at each timepoint relative to the time of DAF.

Of course, estimates of the normalized learning rate become highly uncertain when correlations with DAF escape are small (*e*.*g*., outside the contingent window). However, there is some correlation with DAF escape outside the contingent window and we wanted to include these regions in our analysis. Thus, to identify regions of unreliability in our estimate of the normalized learning rate, we computed the reliability ratio at each timepoint (an estimate of how much larger the correlation is than its noise), as follows:

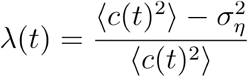

where *c*(*t*) is the correlation with DAF at each timepoint and *σ*_*η*_ is an estimate of the noise in *c*(*t*). For timepoints with *λ* (*t*) < 0.5, we imposed a large fixed uncertainty in the estimate of 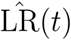. These are plotted in Fig. 5d as blue shaded regions.

Finally, we compute the algorithmic eligibility trace, *e*_*a*_(*t*), from the normalized learning rate using Wiener deconvolution. 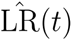 was modeled as

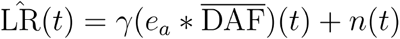

Where 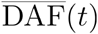 is the DAF profile (the fraction of motifs where DAF was present at each timepoint), *n*(*t*) is an estimate of the measurement noise, and *γ* is a scalar factor. We took *n*(*t*) to be white, and modeled the autocorrelation of *e*_*a*_(*t*) as that of a Gaussian function with width given by our estimate of the learning kernel. We then chose the smallest value of *γ* such that the sum of squared residuals between the measured 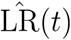 and our estimate of 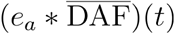 was comparable to an estimate of the noise variance.

#### Premotor latency estimation

Instead of attempting to estimate the effect of a single LMAN neuron on song, we leveraged our observation that nearby LMAN neurons exhibit correlated activity during singing and sought to estimate the effect of the summed activity of ensembles of correlated neurons on the song. To identify LMAN ensembles, we computed pairwise, Spearman correlations between the spike counts of LMAN neurons in 10 ms bins during song. After setting correlation values less than 0.05 to 0, these pairwise correlations were used to define weighted edges in a graph of all LMAN units in a given recording. We then used the Louvain algorithm to identify communities of correlated LMAN neurons. Only ensembles containing 5 or more neurons were used in the following analyses (Extended Data Fig. 7a).

We then estimated the effect on song of changes in the activity of each ensemble as follows. First, we picked a 10 ms long window in the song. All motifs in the dataset were then sorted by the number of spikes fired by the chosen LMAN ensemble within this 10 ms window. The mean was taken over all song spectrograms in the top 50th percentile to produce an average “high activity” spectrogram, *S*_high_(*t, f*). Similarly, the mean was taken over all song spectrograms in the bottom 50th percentile to produce an average “low activity” spectrogram, *S*_low_(*t, f*). The effect of activity at the chosen time was defined as the difference between these two spectrograms,

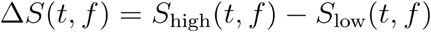

(Extended Data Fig 7b,c). The difference magnitude was then computed as the log of the Euclidean norm of the difference spectrogram over the frequency axis,

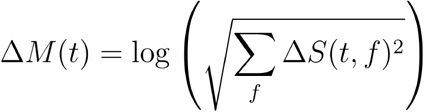

To account for differences in spectral variability at different times in song, this difference magnitude was z-scored using a generated distribution of random difference magnitudes (n=1000).

This process was repeated for all LMAN ensembles and a series of overlapping, 10 ms bins with 5 ms steps between them (Extended Data Fig. 7d). All difference magnitudes were then shifted so that the bins of LMAN ensemble activity were aligned and the mean was taken to produce an average magnitude curve.

To account for longer timescale correlations between LMAN activity and song, this entire process was repeated with motif spectrograms shifted by one motif relative to their associated LMAN activity, in either direction. The average of the +1 and −1 shifted curves were subtracted from the original curve to produce the final result. 95% confidence intervals were bootstrapped (n=1000) by resampling with replacement the different magnitudes corresponding to different activity windows.

The combined effect-on-song was computed as the mean of the curves for each bird. Note that to better quantify the value of the premotor latency, the 3 birds with the most visible effect of LMAN activity on song, as based on their difference spectrograms, were used. Note that in the case of one bird, Yellow33, two variants of the motif were sung, where the second of four syllables took two different forms. For this bird, motifs were categorized into variant 1 and variant 2 and analyzed separately, before combining in the final curve.

The premotor latency was then estimated by first finding the peak of the average difference magnitude in the −50 ms to 50 ms time window. The centroid across the region of the curve about the half-height of this point was then taken as the estimated premotor latency. 95% confidence intervals were found by bootstrapping (n=1000).

#### pCAF learning rate analysis

Measured pitches in the contingent window were first corrected for baseline, circadian fluctuations in pitch across each day. A Gaussian process (GP) model was fit to the measured pitches at their respective time of day, across all days of experiments for a given bird. Times of day were jittered uniformly by up to 30 minutes to prevent overfitting. The pitch predicted by the GP at each time of day was then subtracted from the measured pitches before any additional analysis.

For each day-long pCAF experiment, learning rates were estimated by performing a repeated median regression with the motif number as the independent variable and the pitch as the dependent variable. Learning rates were then normalized by the correlation between short-term pitch changes and DAF, given by the Spearman correlation coefficient between the differences between pitches in consecutive motifs, and the difference in a binary variable representing whether DAF was or was not played.

The predicted curve of learning rate versus DAF delay was generated by convolving a 100 ms square pulse, with its onset at zero (representing the 100 ms acoustic noise burst), with the estimated algorithmic eligibility trace (Fig. 5e). Confidence intervals were bootstrapped based on uncertainty in the algorithmic eligibility trace. This gave a prediction for the overall shape of learning rate versus DAF delay, but said nothing about the absolute magnitudes of learning. Thus, for each bird we fit a unique scale factor to the predicted learning curve using ordinary least squares (OLS) regression:

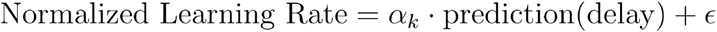

where *α*_*k*_ is the scale constant for each bird. Note that the measured normalized learning rates at delays of ~305ms were not used in fitting the model, but were instead used to measure the variance of the noise in the learning rate measurements. This is justified by prior findings that pCAF does not drive learning for DAF delays greater than 100ms^29^. The noise estimate was used in conjunction with the measured residuals of the model to compute a reduced chi-squared statistic. To facilitate plotting data from all birds together, the learning rates plotted in Fig. 5g are the normalized learning rates, divided by the fitted scale factor, *α*_*k*_, for each bird.

## Statistics and reproducibility

Experiments and analyses were not performed blind to the conditions of the experiments. Statistical methods were not used to predetermine sample sizes, but our sample sizes are similar to those reported in previous studies. Statistical tests are described in the Methods section for each analysis. Unless otherwise stated, α=0.05 was used as the threshold for significance and bootstraps were performed with n=1000 replicates. Data were analysed using MATLAB (MathWorks).

## Data and code availability

All preprocessed data and code needed to reproduce the analysis and generate the figures in this paper are available at https://github.com/FeeLab/ncaf_paper_code. The raw datasets generated and analysed during the current study are not publicly available due to their size, but are available from the corresponding author on reasonable request.

### Acknowledgements

We thank Andrew Bahle, Mark Goldman and Simon Bauerle for critical comments on the manuscript. Funding was provided by the Simons Collaboration on the Global Brain. JRS acknowledges funding through the Harold and Ruth Newman Family Hertz Graduate Fellowship. JMW acknowledges funding through the American Australian Association Graduate Education Scholarship, the Quad Fellowship, and the MathWorks Fellowship.

## Author Information

### Contributions

The study was conceived and designed by JRS, JMW and MSF. Behavioral pitch-learning data were collected by ANDP. Other experimental data were collected by JRS and JMW. Data were analysed by JRS, JMW and MSF. JRS, JMW and MSF wrote the manuscript. All authors contributed to reviewing and editing the manuscript.

---

**Extended Data Figure 1:**
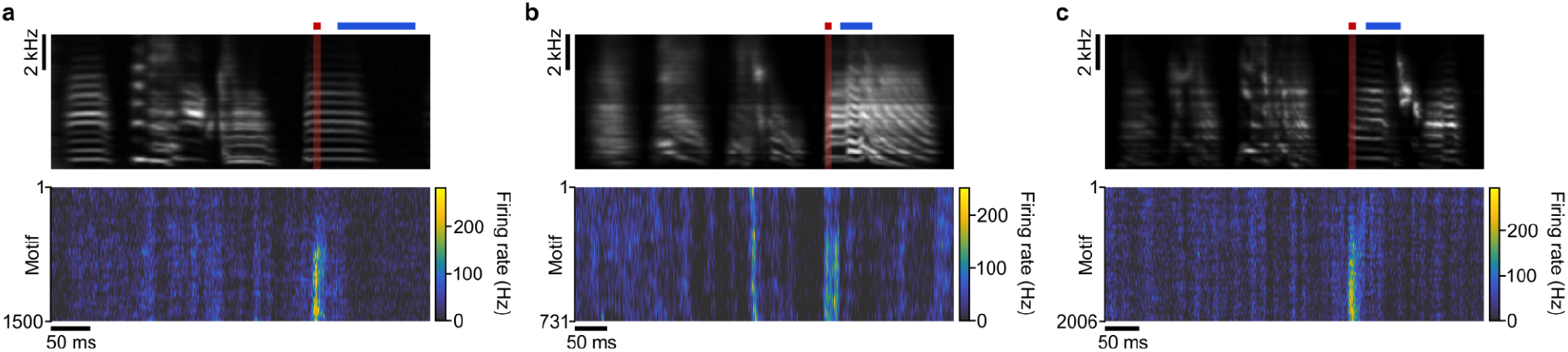
Additional nCAF example rasters. **a**-**c** spectrograms (top panels) and rasters (bottom panels) of neural activity in a targeted single neuron across three nCAF experiments in three different birds, plotted as in Fig. 1c. DAF was delivered when neural activity was lower than average within the contingent window.

**Extended Data Figure 2:**
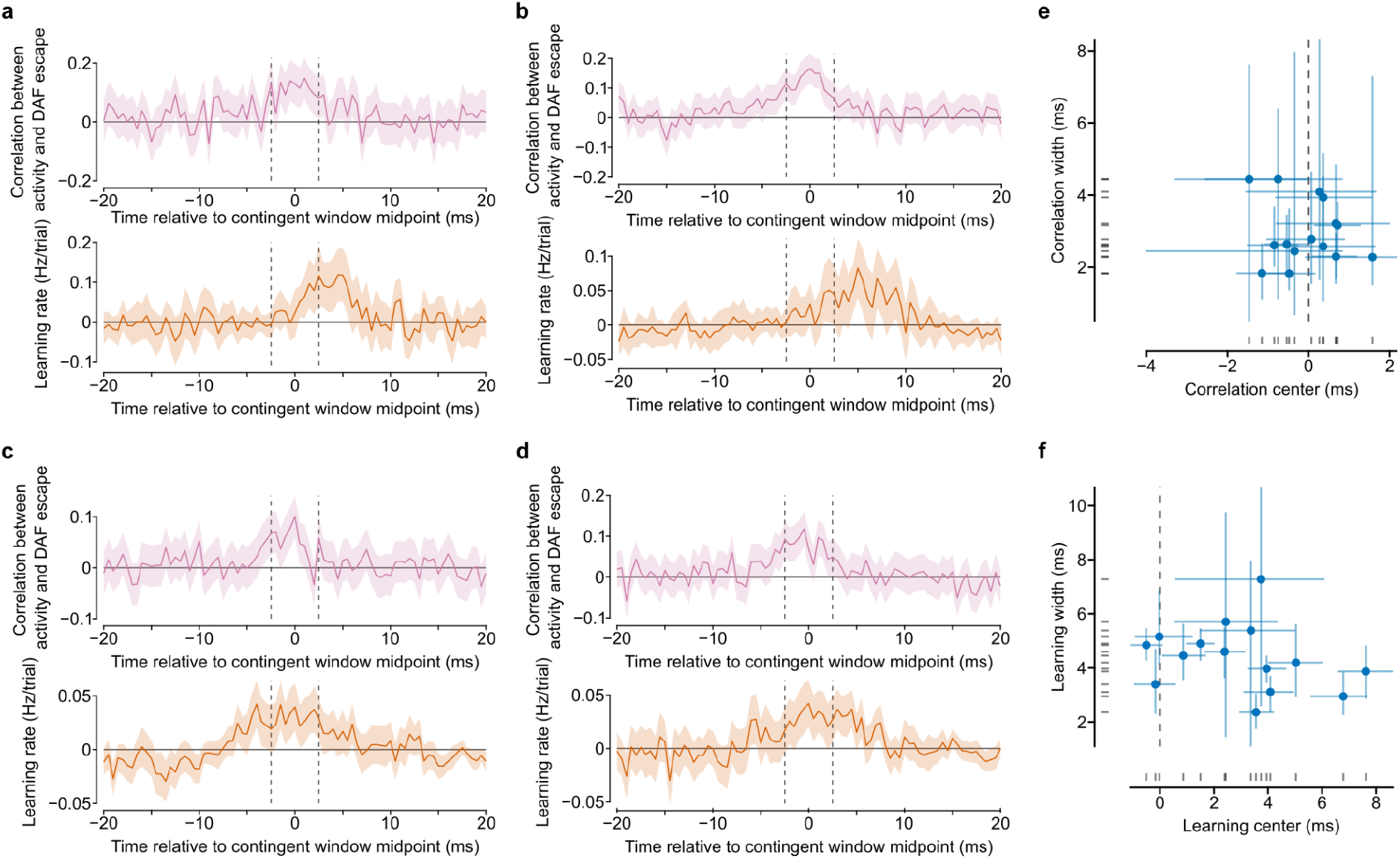
Additional examples of correlation and learning profiles used to measure the temporal precision of nCAF learning. **a**-**d**, Correlation profile (top panel) and learning profile (bottom panel) for 4 additional targeted units, plotted as in Figure 2a,b. **e**, Offset and width of Gaussian fits to the correlation profile of each unit (error bars are 95% CI). **f**, As in panel e but for fits to learning profiles for each unit.Timing precision across neurons

**Extended Data Figure 3:**
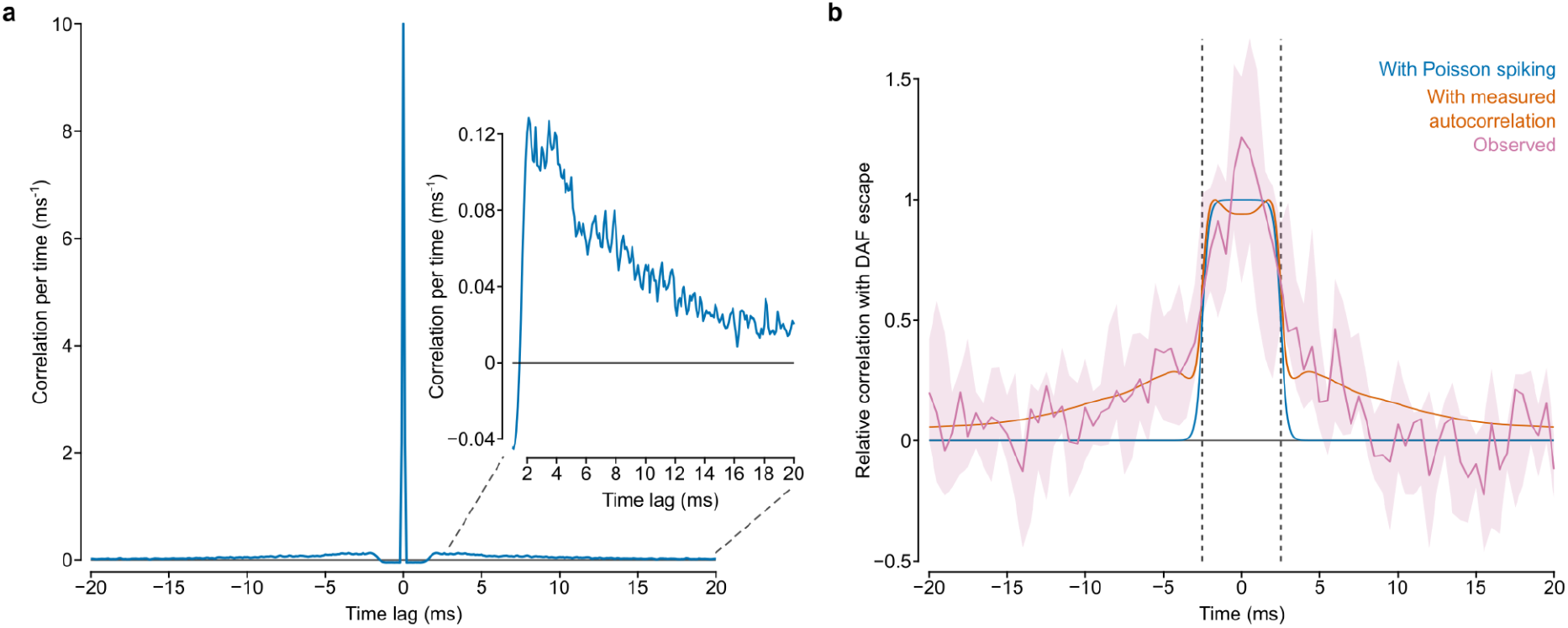
Effects of LMAN autocorrelations on correlation profiles imposed by nCAF. We examine LMAN spiketrain autocorrelations to determine whether these contribute to the fact that the correlation profiles we observe are wider than the contingent window. If LMAN spiketrains obeyed a Poisson process, then they would exhibit no temporal autocorrelations and each unit’s correlation profile would simply be given by the average location of the contingent window. However, because LMAN neurons generate bursts and have slower firing rate fluctuations, spike counts within the contingent window are correlated with spike counts just outside. This leads to a broadening of the correlation profile. **a**, Average, mean-subtracted autocorrelation for the LMAN single units in Fig. 3b, calculated using 100 μs windows. Note the presence of refractory period at lags < 2ms, and positive sidelobes at longer times (>2ms). **b**, Comparison between the predicted correlation profile (blue) for a neuron with a Poisson spiketrain (simply the average contingent window location), the predicted correlation profile (orange) for a neuron with observed LMAN spiketrain autocorrelations (computed by convolving the average contingent window location with the measured LMAN spiketrain autocorrelation), and the median correlation profile (pink) observed across targeted units (shading is 95% CI; vertical dashed lines indicate contingent window edges). Note that the width of the observed correlation profile (pink) is consistent with the width of the predicted correlation profile (orange), based on LMAN spiketrain autocorrelations. While there is some broadening of the correlation profile relative to the shape of the contingent window, the full-width half-maximum is essentially unchanged relative to that of the correlation profile of a Poisson LMAN spiketrain. Thus, despite LMAN spiketrain autocorrelations, the learning circuit has the same narrow temporal resolution as if the LMAN spiketrain were a Poisson process.

**Extended Data Figure 4:**
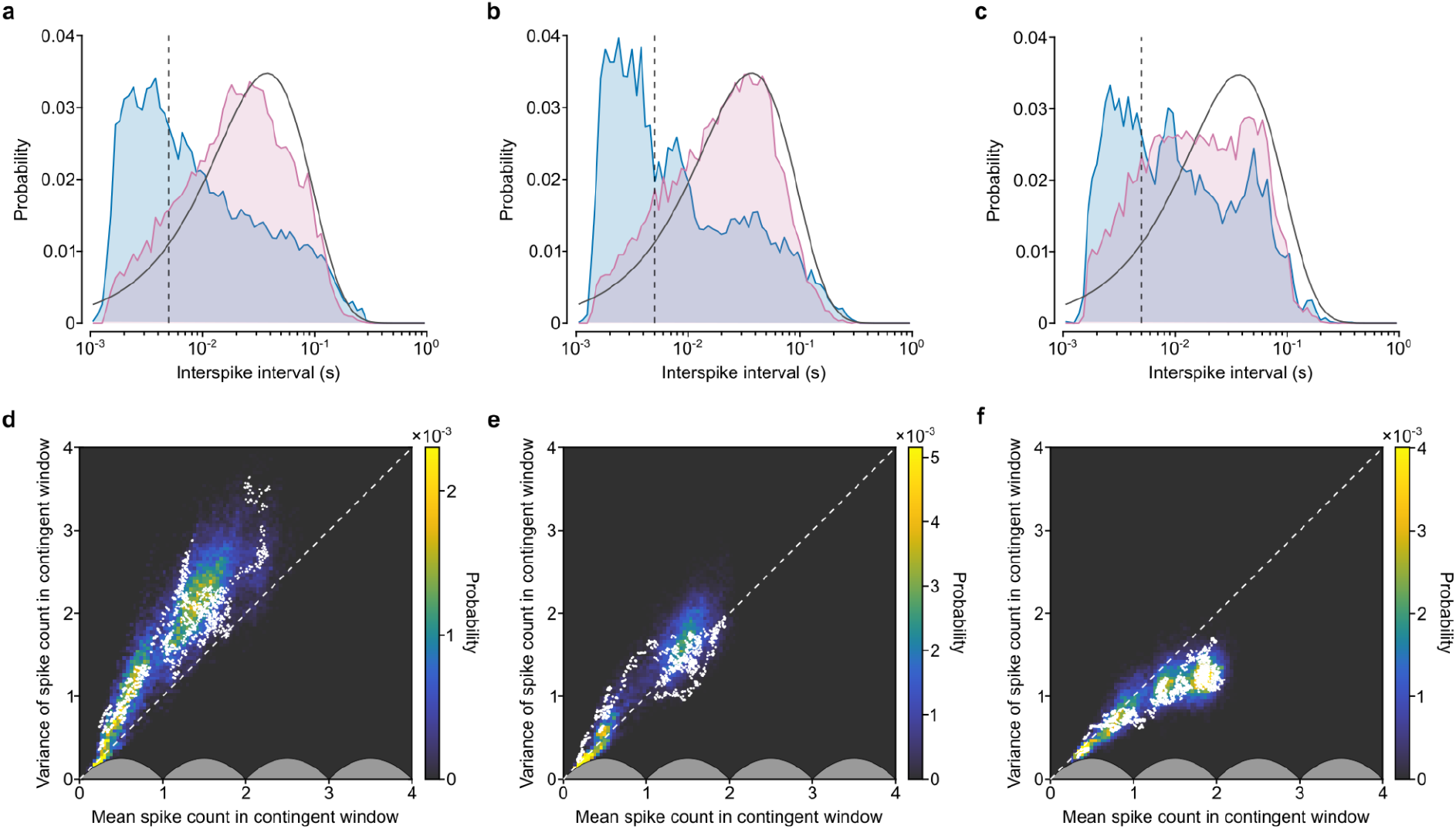
Spike statistics across neurons. **a**-**c**, Interspike interval distributions (not including the contingent window) for three neurons in Fig. 3b (blue), simulated inhomogeneous Poisson processes with the same rate parameter modulation as each neuron (pink), and simulated homogenous Poisson processes with the same average rate parameter (gray outline). **d**-**f**, White points: mean and variance of the number of spikes in the contingent window for three single neurons, computed across a sliding window of 100 consecutive motifs. Heatmaps: distributions of observed spike statistics in the contingent window for simulated models of best fit that incorporate both bursting and Poisson firing.

**Extended Data Figure 5:**
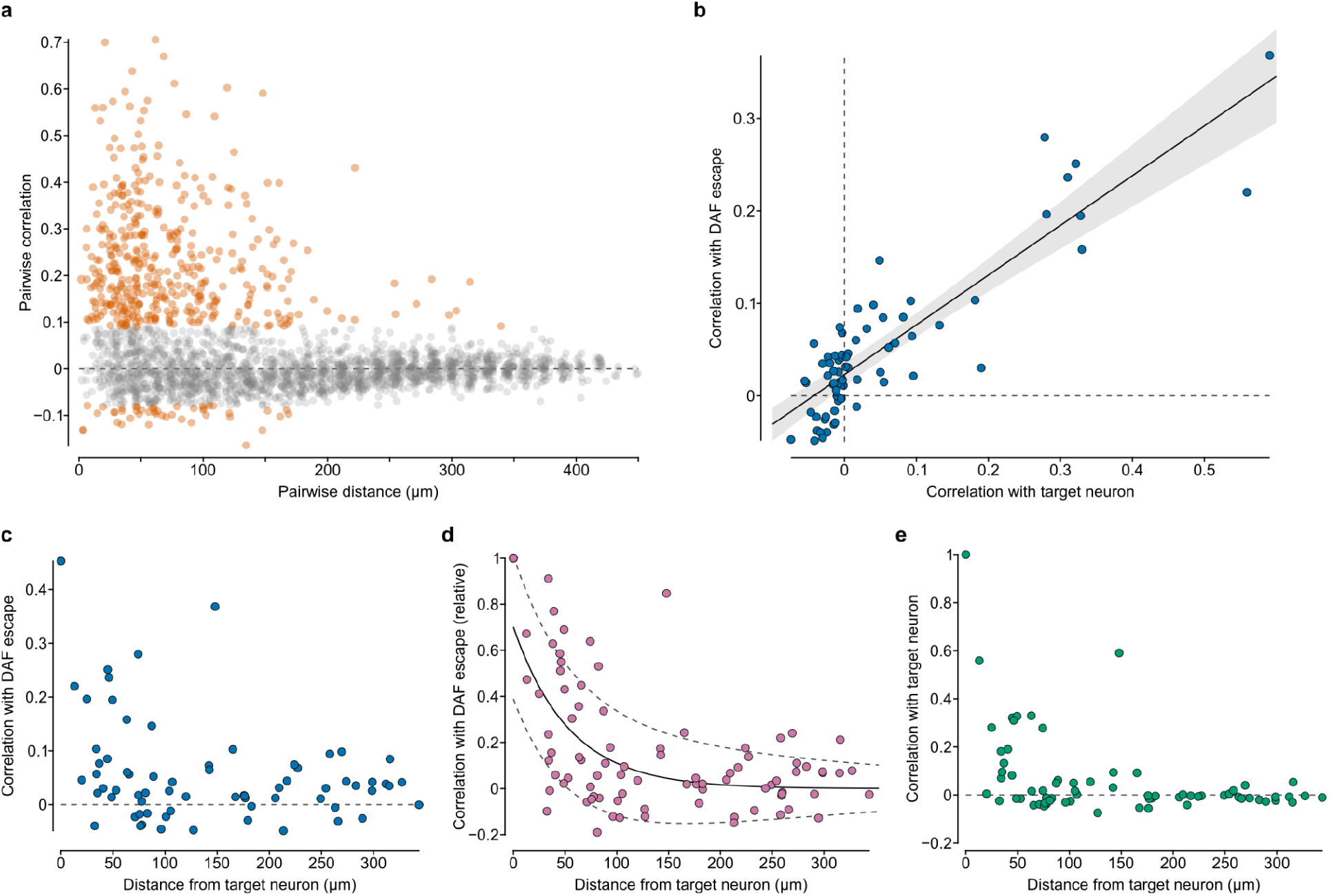
Spatial learning statistics. **a**, Correlations between pairs of LMAN neurons as a function of pairwise distance (n=69 neurons). Pairs with statistically significant correlations (95% confidence level compared to shuffled data, Bonferroni correction) are plotted in orange. **b**, Correlation with DAF escape versus correlation with the target neuron for untargeted single neurons in the dataset in Fig. 4. A line of best fit is plotted in black (*β* = 0.54 ± 0.07, 95% CI). **c**, Correlation with DAF escape versus distance from the target neuron for the same dataset. **d**, Correlation between neural activity and DAF escape in the same dataset, normalized in each experiment by the correlation of the target neuron with DAF. Plotted line and 95% CI are for a parameterized distribution fit to the data (see Methods). **e**, Correlation with the target neuron versus distance from the target neuron for the same dataset. (Same data as panel a, but only showing pairs that include the target neuron.

**Extended Data Figure 6:**
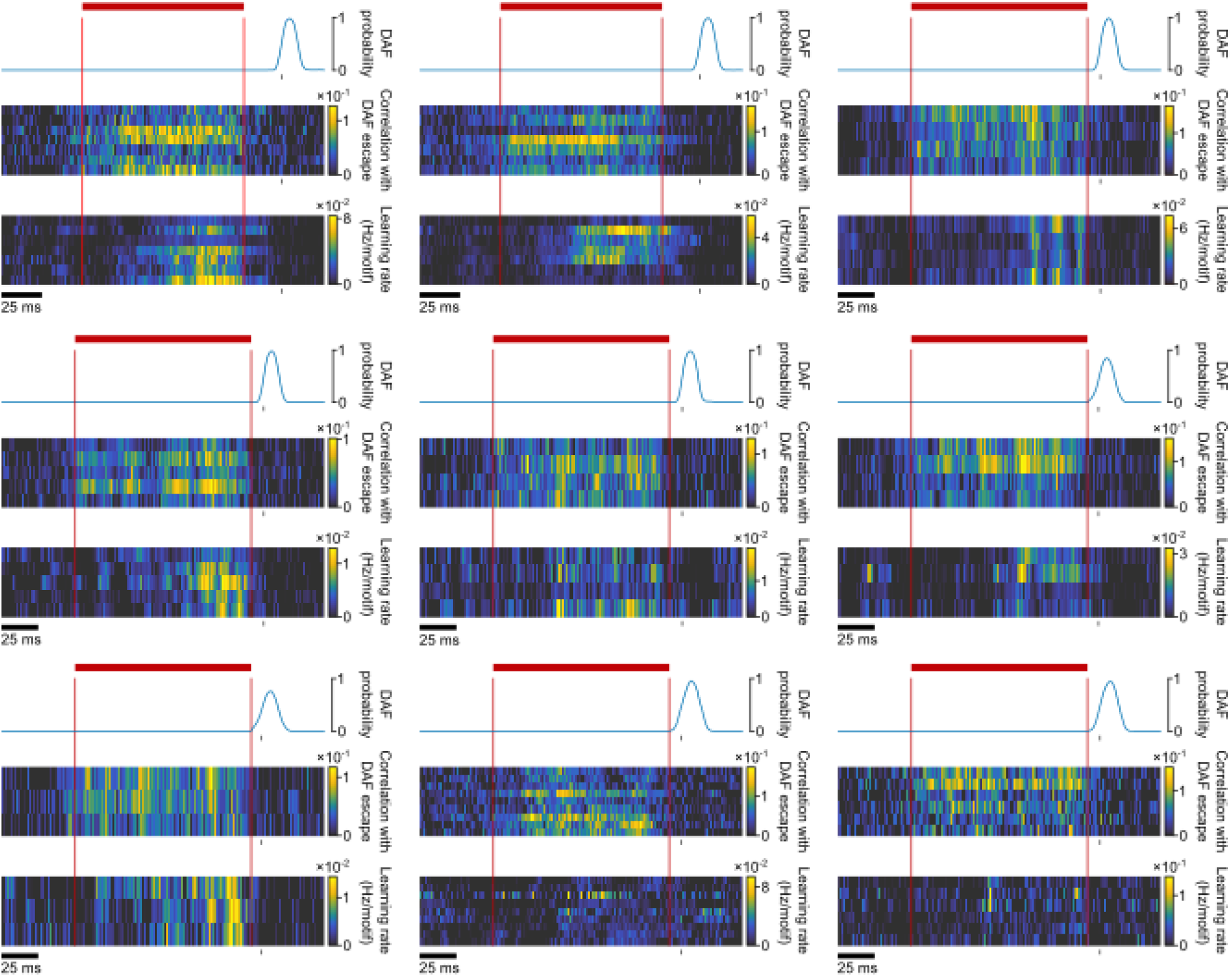
Additional experiments and birds for eligibility trace measurements (Related to Figure 5). Top panels: DAF profile, plotted as the probability that a given timepoint overlaps with DAF on a given motif. Middle panel: correlation between neural activity and DAF for a range of units. Bottom panel: learning rate for the same set of units (edges of the contingent window plotted in red; gray ticks represented time of DAF onset).

**Extended Data Figure 7:**
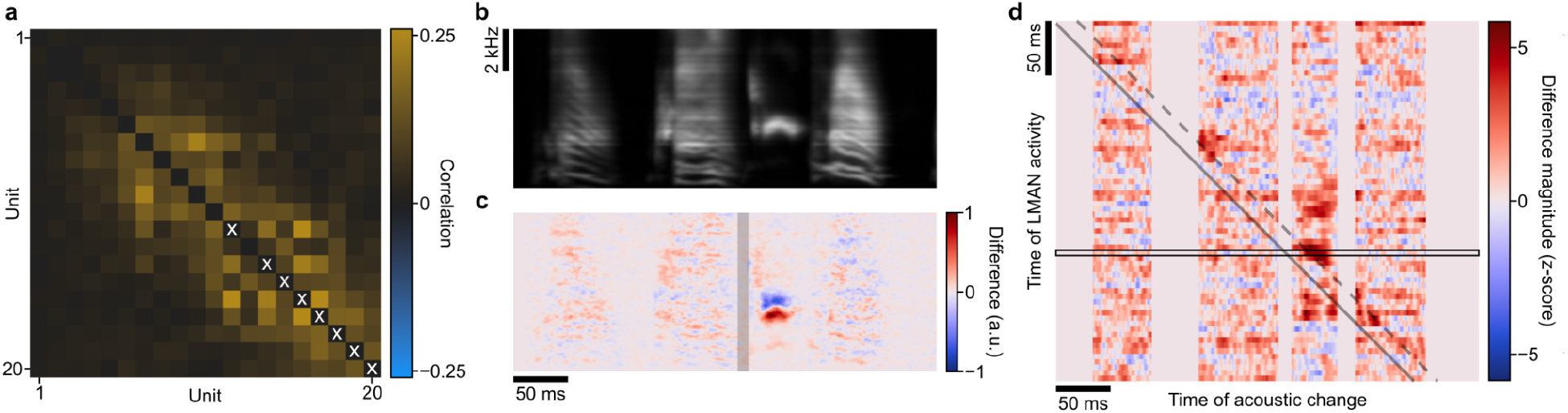
Relationship between LMAN activity fluctuations and acoustic output. **a**, Spearman correlation between neural activity for pairs of LMAN units in an example bird, sorted by depth along the Neuropixels shank. Units were observed to form correlated ensembles. One such ensemble (marked here with white crosses) was used for the analyses in this figure (see Methods). Song renditions sung across the day were split into two groups, based on whether the summed activity of the chosen ensemble within a narrow, 10 ms window was above or below average. The difference was taken between the means of the song spectrograms in each group to give a difference spectrogram, corresponding to activity changes in the chosen window. **b**, Mean song spectrogram sung across the day. **c**, Difference spectrogram for the activity window shaded in gray. This was repeated for all 10 ms windows in the song. The magnitude of each difference spectrogram over time was then taken and z-scored relative to a bootstrapped distribution of random difference spectrogram magnitudes. **d**, Matrix of z-scored spectral difference magnitudes computed based on LMAN activity across a range of 10 ms windows. Each row is the magnitude of a single difference spectrogram contingent on activity of the chosen ensemble in a different 10 ms window. Black rectangle corresponds to the difference spectrogram in panel c. Solid gray diagonal line marks the center of the LMAN ensemble activity window used to create each row. The projection of this matrix along this diagonal was used to contribute to the plot in Fig. 5f. Dashed gray off-diagonal line indicates the time of the estimated premotor latency, 21.6 ms. Note the presence of regions of large spectral difference along this line.

