## Supplementary Note for "Millisecond-scale, single-neuron credit assignment in a songbird"

### Supplementary Note:
### Algorithmic and Synaptic Eligibility Traces

Eligibility traces are a solution to the temporal credit assignment problem. They serve as transient records that allow outcomes to be associated with the states, actions, or both that preceded them[1–3]. Here, we restrict our discussion to eligibility traces in the context of biological neural networks. In this context, an eligibility trace is typically conceptualized as a molecular tag associated with a synapse. When that synapse is *active*, an eligibility trace is established. The eligibility trace decays over time, but is longer-lived than the event that established it. When there is subsequent input from a neuromodulatory third factor (*e.g.*, dopamine), the strength of each synapse is updated according to the value of its eligibility trace.

#### Synaptic eligibility traces in the song system

Consider a single synapse from an HVC neuron onto a medium spiny neuron (MSN) in Area X, which also receives input from a neuron in LMAN. In our model, coincident input from the HVC and LMAN neurons establishes a synaptic eligibility trace specifically at the HVC→MSN synapse[4]. The weight of the synapse is then updated based on the temporal overlap between the synaptic eligibility trace and subsequent dopaminergic reward-prediction error (RPE) signals from the ventral tegmental area (VTA). Suppose the HVC input to the synapse is active at time $t = 0$. Then, we can write the update to the weight of this synapse on motif $n$, $\Delta w_n$, as follows:

$$\Delta w_n = L_n \cdot \int_{-\infty}^{\infty} e_s(t) \cdot \mathrm{DA}_n(t)\, dt \tag{1}$$

where $L_n$ is a binary variable representing whether there was coincident LMAN input on motif $n$, $e_s(t)$ is the time course of the synaptic eligibility trace, and $\mathrm{DA}_n(t)$ is the time course of the dopamine transient on motif $n$.

In our nCAF experiment, let us assume that the dopamine signal on each motif is dominated by the contribution of DAF playback, which occurs on 50% of motifs. In this case, the reward-prediction errors on motifs with and without DAF should be equal in magnitude and opposite in sign (so that the expected RPE is zero). Then, we can write that

$$\mathrm{DA}_n(t) = d_n \cdot \overline{\mathrm{DA}}(t) \tag{2}$$

where $d_n \in \{-1, 1\}$ is a variable representing whether DAF was played (+1) or withheld (-1) on motif $n$, and $\overline{\text{DA}}(t)$ is the shape of the average dopamine transient in response to DAF. Substituting Eq. (2) into Eq. (1), we get that

$$\Delta w_n = L_n \cdot d_n \cdot \int_{-\infty}^{\infty} e_s(t) \cdot \overline{\text{DA}}(t)\, dt. \tag{3}$$

The expected value of this weight update is then

$$\mathbb{E}[\Delta w_n] = \mathbb{E}[L_n \cdot d_n] \cdot \int_{-\infty}^{\infty} e_s(t) \cdot \overline{\text{DA}}(t)\, dt.$$

As DAF is played on 50% of motifs, $\mathbb{E}[d_n] = 0$. Thus, $\mathbb{E}[L_n \cdot d_n]$ is proportional to the correlation between $L_n$ and $d_n$. Therefore,

$$\mathbb{E}[\Delta w_n] \propto \rho \cdot \int_{-\infty}^{\infty} e_s(t) \cdot \overline{\text{DA}}(t)\, dt$$

where $\rho$ is the correlation between DAF and LMAN activity at the time of the HVC input to the synapse.

So far, we have only considered the weight updates to a single HVC→MSN synapse, with HVC input at time $t = 0$. It is straightforward to generalize this to give the expected weight update of any HVC→MSN synapse, with HVC input at time $t = \tau$. We simply need to shift the eligibility trace by $\tau$ and use the correlation between DAF and LMAN activity at time $\tau$:

$$\mathbb{E}[\Delta w_n](\tau) \propto \rho(\tau) \cdot \int_{-\infty}^{\infty} e_s(t - \tau) \cdot \overline{\text{DA}}(t)\, dt.$$

Notably, the integral in this equation can be expressed as a convolution,

$$\mathbb{E}[\Delta w_n](\tau) = \rho(\tau) \cdot \left[e_s(-t) * \overline{\text{DA}}(t)\right]_{t=\tau}$$

where $*$ is the convolution operator. It can also be expressed as a cross-correlation,

$$\mathbb{E}[\Delta w_n](\tau) = \rho(\tau) \cdot \left[e_s \star \overline{\text{DA}}\right](\tau)$$

where $\star$ is the cross-correlation operator.

Up to this point, many of the variables in our equations are experimentally inaccessible. We now relate this result to observable quantities in our experiments. Recall that in our model, strengthening of an HVC→MSN synapse causes the MSN to fire at the time corresponding to the HVC input, in turn driving activity in the corresponding LMAN neuron at that same time. Let us assume that the driven LMAN activity is proportional to the weight of the synapse. Then, if we define $\text{LR}(\tau)$ as the learning rate (the average change in firing rate per motif) of a given LMAN neuron at time $t = \tau$ in the motif, we can write that

$$\text{LR}(\tau) \propto \rho(\tau) \cdot \left[e_s \star \overline{\text{DA}}\right](\tau)$$

or, if we define $\widehat{\mathrm{LR}}(\tau)$ as the learning rate at time $\tau$, divided by the correlation between LMAN activity and DAF at time $\tau$, then

$$\widehat{\mathrm{LR}}(\tau) \propto \left[e_s \star \overline{\mathrm{DA}}\right](\tau) \tag{4}$$

Unlike direct synaptic weight measurements, the learning rate of LMAN neurons is experimentally accessible. However, we do not have direct access to the average dopamine response to DAF. We do, however, have access to the average time course of the DAF itself. Let us assume that there is a linear, time-invariant mapping from *true* reward-prediction errors to dopamine transients. Then, we can say that

$$\mathrm{DA}(t) = [K_{\mathrm{DA}} * \mathrm{RPE}](t) \tag{5}$$

where $\mathrm{RPE}(t)$ is the reward-prediction error at each point in time and $K_{\mathrm{DA}}(t)$ is some underlying dopamine kernel. Then, substituting Eq. (5) into Eq. (4),

$$\widehat{\mathrm{LR}}(\tau) \propto \left[e_s \star \left[K_{\mathrm{DA}} * \overline{\mathrm{RPE}}\right]\right](\tau)$$

where $\overline{\mathrm{RPE}}(t)$ is the RPE on motifs where DAF is played. This can be rearranged to give

$$\widehat{\mathrm{LR}}(\tau) \propto \left[[K_{\mathrm{DA}} \star e_s] \star \overline{\mathrm{RPE}}\right](\tau)$$

If we define the **algorithmic eligibility trace** as

$$e_a(t) = [K_{\mathrm{DA}} \star e_s](t)$$

then

$$\widehat{\mathrm{LR}}(\tau) \propto \left[e_a \star \overline{\mathrm{RPE}}\right](\tau)$$

If we note that in nCAF, the time course of the *true* RPE on motifs where DAF is played is proportional to the average time course of the DAF itself, $\overline{\mathrm{DAF}}(t)$, then we find that

$$\widehat{\mathrm{LR}}(\tau) \propto \left[e_a \star \overline{\mathrm{DAF}}\right](\tau) \tag{6}$$

Now, we have an expression for the normalized learning rate of an LMAN neuron at each point in time in terms of the average DAF time course, which we can measure, and a single unknown function: the algorithmic eligibility trace. Critically, this means that the algorithmic eligibility trace is recoverable by deconvolution, without knowing the dopamine kernel, $K_{\mathrm{DA}}$.

#### Relating the algorithmic and synaptic eligibility traces

Even if we do not know the exact form of the dopamine kernel, $K_{\mathrm{DA}}(t)$, we can use an estimate of the algorithmic eligibility trace to make inferences about the synaptic eligibility trace. In particular, it is a property of convolution that if we have two functions, $x(t)$ and $y(t)$, which extend over the domains $[t_x^i, t_x^f]$ and $[t_y^i, t_y^f]$, respectively, then the convolution of $x(t)$ and $y(t)$ will extend over the domain $[t_x^i + t_y^i, t_x^f + t_y^f]$. Recalling the relationship between convolution and cross-correlation and applying this result, we can say that

$$t_{e_a}^i = t_{e_s}^i - t_{K_{\mathrm{DA}}}^f$$
$$t_{e_a}^f = t_{e_s}^f - t_{K_{\mathrm{DA}}}^i$$

or, equivalently,

$$t_{e_s}^i = t_{e_a}^i + t_{K_{\mathrm{DA}}}^f$$
$$t_{e_s}^f = t_{e_a}^f + t_{K_{\mathrm{DA}}}^i$$

Thus, knowing the onsets and offsets of the algorithmic eligibility trace and dopamine kernel allows us to infer the onset and offset of the synaptic eligibility trace. In the main text, we use the known onset of the dopamine kernel and our estimate of the offset of the algorithmic eligibility trace to predict the offset of the synaptic eligibility trace.
